# Transcriptome analysis of *Pseudomonas aeruginosa* biofilm infection in an *ex vivo* pig model of the cystic fibrosis lung

**DOI:** 10.1101/2021.07.23.453509

**Authors:** Niamh E. Harrington, Jenny L. Littler, Freya Harrison

## Abstract

*Pseudomonas aeruginosa* is the predominant cause of chronic biofilm infections that form in the lungs of people with cystic fibrosis (CF). These infections are highly resistant to antibiotics and persist for years in the respiratory tract. One of the main research challenges is that current laboratory models do not accurately replicate key aspects of a chronic *P. aeruginosa* biofilm infection, highlighted by previous RNA-sequencing studies. We compared the *P. aeruginosa* PA14 transcriptome in an *ex vivo* pig lung (EVPL) model of CF and a well-studied synthetic cystic fibrosis sputum medium (SCFM). *P. aeruginosa* was grown in the EVPL model for 1, 2 and 7 days, and *in vitro* in SCFM for 1 and 2 days. The RNA was extracted and sequenced at each time point. Our findings demonstrate that expression of antimicrobial resistance genes was cued by growth in the EVPL model, highlighting the importance of growth environment in determining accurate resistance profiles. The EVPL model created two distinct growth environments: tissue-associated biofilm and the SCFM surrounding tissue, each of which cued a transcriptome distinct from that seen in SCFM *in vitro*. The expression of quorum sensing associated genes in the EVPL tissue-associated biofilm at 48 h relative to *in vitro* SCFM was found to be similar to CF sputum versus *in vitro* conditions. Hence, the EVPL model can replicate key aspects of *in vivo* biofilm infection that are missing from other current models and provides a more accurate *P. aeruginosa* growth environment for determining antimicrobial resistance.

**Significance Statement:** *Pseudomonas aeruginosa* lung infections that affect people with cystic fibrosis are a well-researched area, yet they continue to evade antimicrobial treatments. The lack of a laboratory model that captures all key aspects of these infections hinders not only research progression but also clinical diagnostics. We used transcriptome analysis to demonstrate how a model using pig lungs can more accurately replicate key characteristics of *P. aeruginosa* lung infection, including mechanisms of antibiotic resistance and infection establishment. Therefore, this model may be used in the future to further understand infection dynamics to develop novel treatments and more accurate treatment plans. This could improve clinical outcomes as well as quality of life for individuals affected by these infections.

## Introduction

The Gram-negative, facultative anaerobe and opportunistic pathogen *Pseudomonas aeruginosa* is the most common cause of chronic biofilm infection in the lungs of people with the genetic condition cystic fibrosis (CF); associated with reduced life quality and increased mortality [1]. The pathogen’s ability to persist for years in the respiratory tract of people with CF is associated with its adaptive mechanisms. Arguably, the most important of these is the switch to a biofilm lifestyle in this environment, conferring protection to the host immune response and high antibiotic tolerance [2]. Biofilm formation creates a spatial organisation that results in complex cell-to-cell interactions and increased infection population heterogeneity [3]. Although these infections are widely regarded as the best described and most studied biofilm infection in medicine, there are no completely effective eradication strategies [4]. There are a number of limitations with the current laboratory models, and these have impacted research progression [5,6]. The lack of an *in vivo*-like growth platform for current susceptibility testing methods to prescribe drugs often results in poor clinical outcomes [7].

Existing infection models of the CF lung include mouse, ferret, and pig models, as well as biofilms formed on beads or epithelial cultures and laboratory sputum media designed to mimic human CF sputum. The variety of different models, each capturing different features of CF lung infection, has led to RNA sequencing (RNA-seq) studies that aim to identify the transcriptomic aspects of *P. aeruginosa* chronic biofilm infection not captured by current work [8,9]. This approach has highlighted the importance of growth environment in determining *P. aeruginosa* traits involved in virulence, persistence and antimicrobial resistance (AMR), key considerations for treating CF lung infections [8–10]. A quantitative computational framework using transcriptomic data for the evaluation of human infection model accuracy has also been developed [5]. This work found that an *in vitro* CF epithelial cell model and a revised version of a specific synthetic cystic fibrosis sputum medium (SCFM2) both resulted in *P. aeruginosa* gene expression more comparable to the *P. aeruginosa* transcriptome from human CF sputum than other models tested. The key aspects of *in vivo* metabolism captured included fatty acid and phospholipid metabolism in the epithelial cell model, and nucleoside and nucleotide metabolism in SCFM2. However, there are aspects of infection that are not reproduced by these two models. Alginate production and quorum sensing (QS)-associated genes were among the genes that caused the biggest distinction between *in vitro* and *in vivo* transcriptomes (both overexpressed *in vitro*) [5]. Both alginate and QS are directly linked to *P. aeruginosa* persistence and virulence [11,12], thus there is a crucial gap in our ability to reproduce CF-like phenotypes in the laboratory.

We have developed an *ex vivo* pig lung (EVPL) model of the CF lung environment that replicates key phenotypic aspects of *P. aeruginosa* chronic biofilm infection [13–16]. We aimed to assess the *P. aeruginosa* transcriptome as biofilm infection establishes in the EVPL model, to determine the extent to which the model accurately replicates *P. aeruginosa* gene expression during human CF infection. We show that the EVPL model creates two environments that are distinct from SCFM *in vitro* growth, and that it cues patterns of *P. aeruginosa* gene expression that are more comparable to those seen in human infection. When the transcriptome of *P. aeruginosa* grown as a biofilm on the EVPL tissue is compared with that of *P. aeruginosa* in SCFM *in vitro*, the changes in expression of key metabolic pathways, quorum sensing-controlled virulence determinants and genes linked to antibiotic resistance are similar to those observed when the *P. aeruginosa* transcriptome *in vivo* in CF infection is compared with the *in vitro* transcriptome.

## Results

The EVPL model comprises two environments: the bronchiolar lung tissue surface (lung) and the SCFM that surrounds the tissue (surrounding SCFM) to mimic the luminal mucus in the human CF lung (see Fig S1). We aimed to determine the changes in the *P. aeruginosa* PA14 transcriptome cued by growth as either tissue-associated biofilm, or in the surrounding SCFM of the EVPL model, compared with SCFM *in vitro.* PA14 was grown in the EVPL model and in SCFM alone for seven days, and comparisons were made at two time points (24 h and 48 h; we could not recover sufficient mRNA from 7-day populations in SCFM *in vitro*). We used these comparisons to address the overall question: does biofilm growth on pig bronchiolar tissue better replicate the *P. aeruginosa* transcriptome observed in human infection than growth in SCFM *in vitro*?

### EVPL tissue surface maintains viable *P. aeruginosa* populations for longer than SCFM *in vitro*

*P. aeruginosa* PA14 was grown on 3 replica tissue sections from each of two independent lungs for 24 h, 48 h or 7 d. RNA was extracted from the lung-associated biofilm and the surrounding SCFM for each sample. RNA was also extracted at the same time points from PA14 grown in SCFM *in vitro*. At 7 days, RNA concentrations sufficient for sequencing could not be consistently obtained from *P. aeruginosa* grown *in vitro* (Table S1). However, sufficient *P. aeruginosa* RNA was obtained from lung tissue-associated biofilm at this time point. We confirmed that PA14 was still viable on the lung tissue at 7 d post-infection (PI) (Fig S2) and did not appear to be visibly ‘stressed’ as seen for *in vitro* cultures (rounding of cells: Fig S3). Hence, RNA from lung tissue-associated biofilms was sequenced from 24 h, 48 h and 7 d PI, but RNA from *in vitro* cultures and the SCFM surrounding lung tissue was sequenced from 24 h and 48 h only.

**Table 1.**
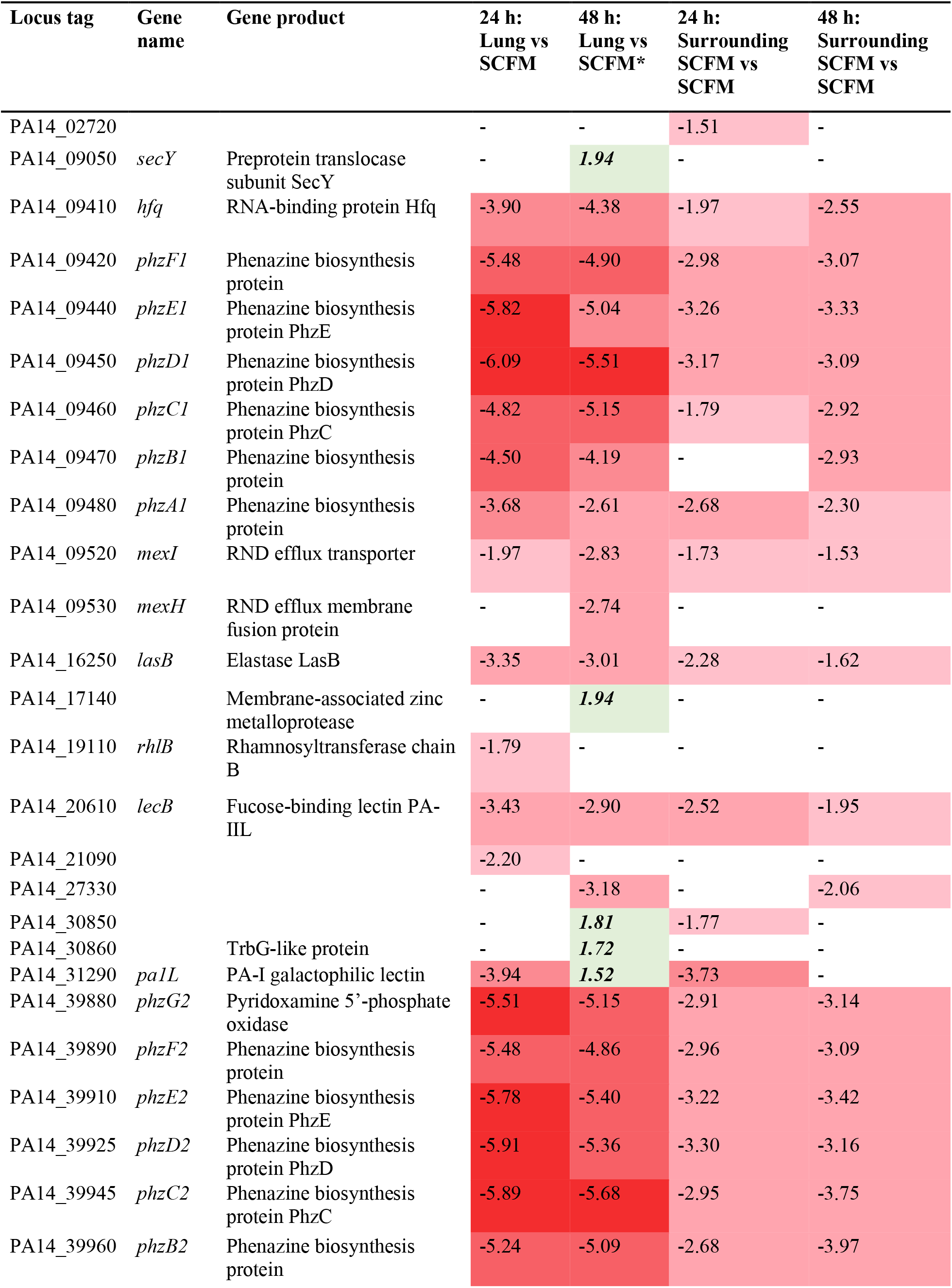
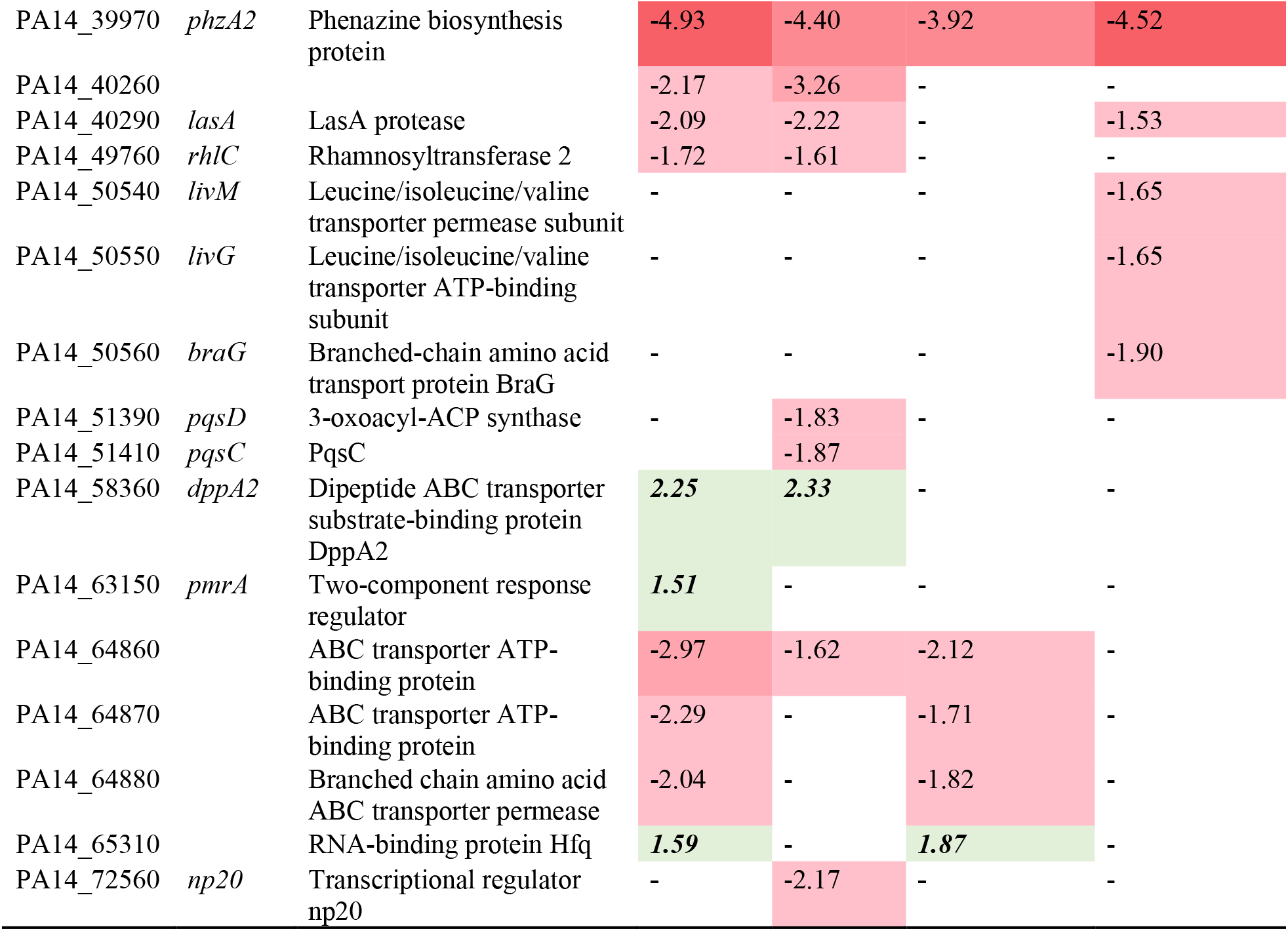
All genes associated with the *Pseudomonas aeruginosa* PA14 KEGG pathway quorum sensing as listed on Pseudomonas.com [24] that were found to be significantly differentially expressed (*P* < 0.05, log_2_ fold change ≥ |1.5|) in at least one contrast. PA14 expression in the two locations of the *ex vivo* pig lung model: the lung tissue-associated biofilm (lung) and the synthetic cystic fibrosis sputum media (SCFM) surrounding the lung tissue (surrounding SCFM), was compared with *in vitro* SCFM (SCFM) growth. Samples were compared at 24 h and 48 h post infection. The log_2_ fold change (lfc) is shown where significant, and contrasts donated with ‘-’ for a gene show it was not significantly differentially expressed. The red fill shows genes that are underexpressed (also standard font) in the EVPL environments versus *in vitro* SCFM, the darker the colour the greater the lfc. The green fill shows genes that are overexpressed in the EVPL environments (bold and italicised font). The * denotes the only contrast where the quorum sensing KEGG pathway was not significantly enriched.

Colony forming units (CFU) were measured from representative repeats for each condition used for RNA-seq, to confirm the presence of PA14 and check for consistent growth between environments (Fig S4). Briefly, recovered CFU was comparable between *in vitro* SCFM and the lung tissue at 24 h and 48 h PI, and this density was maintained at 7 d PI on the lung tissue. The CFU ml^−1^ recovered from the surrounding SCFM was approximately one log_10_ higher than that recovered from either the tissue-associated biofilm or *in vitro* cultures at both 24 h and 48 h PI. However, across all environments and time points, the PA14 CFU was consistent with the high densities and variability of CFU that are recovered from people with CF (Fig S4): chronic *P. aeruginosa* lung infection has an expected bacterial load of 10^8^ - 10^10^ ml^−1^ [17].

### *P. aeruginosa* transcriptome analysis reveals distinct niches in the EVPL model compared with *in vitro* SCFM, and distinct changes in transcriptome over time

Following initial data preparation, all reads from EVPL environments (lung and surrounding SCFM) were aligned to the pig genome to exclude any contaminating porcine RNA. There were ≤ 1% of reads aligned to the pig genome across all samples (median 0.09%, see Table S2). Subsequently, remaining EVPL model reads and *in vitro* SCFM reads were successfully aligned to the *P. aeruginosa* PA14 genome (median 98.9%, Table S2) proving the bacterial RNA extracted was predominantly *P. aeruginosa* PA14. This confirmed that there were few metabolically active endogenous microbes present in the EVPL tissue.

Initial principal component analysis (PCA), considering 5829 genes, demonstrated that the different environments in the EVPL model resulted in distinct *P. aeruginosa* PA14 expression profiles (Fig 1A). This suggests that the environmental cues the bacteria encounter may differ between lung tissue-associated biofilm and the airway mucus, resulting in two distinct infection populations. There was also a difference in how expression changed over time when PA14 was grown in the EVPL model or in SCFM *in vitro*. As shown in Figure 1A, the shift in the first two principal components from 24 h to 48 h (and 7 d in the lung-associated biofilm) followed opposite directions for *ex vivo* growth compared with SCFM *in vitro*. This highlights that as well as distinct overall differences in *P. aeruginosa* gene expression between the three environments, there was also a difference in how the expression profile changed over time as the populations established. Subsequent Pearson’s correlation coefficient analysis (*r* > 0.9 for all comparisons) and hierarchical clustering (Fig 1B) showed that the transcriptome of PA14 grown on lung tissue at the two later time points (48 h and 7 d) was distinct from PA14 in the surrounding SCFM at 24 h, and growth in SCFM *in vitro* at both time points (Fig 1B).

**Figure 1.**
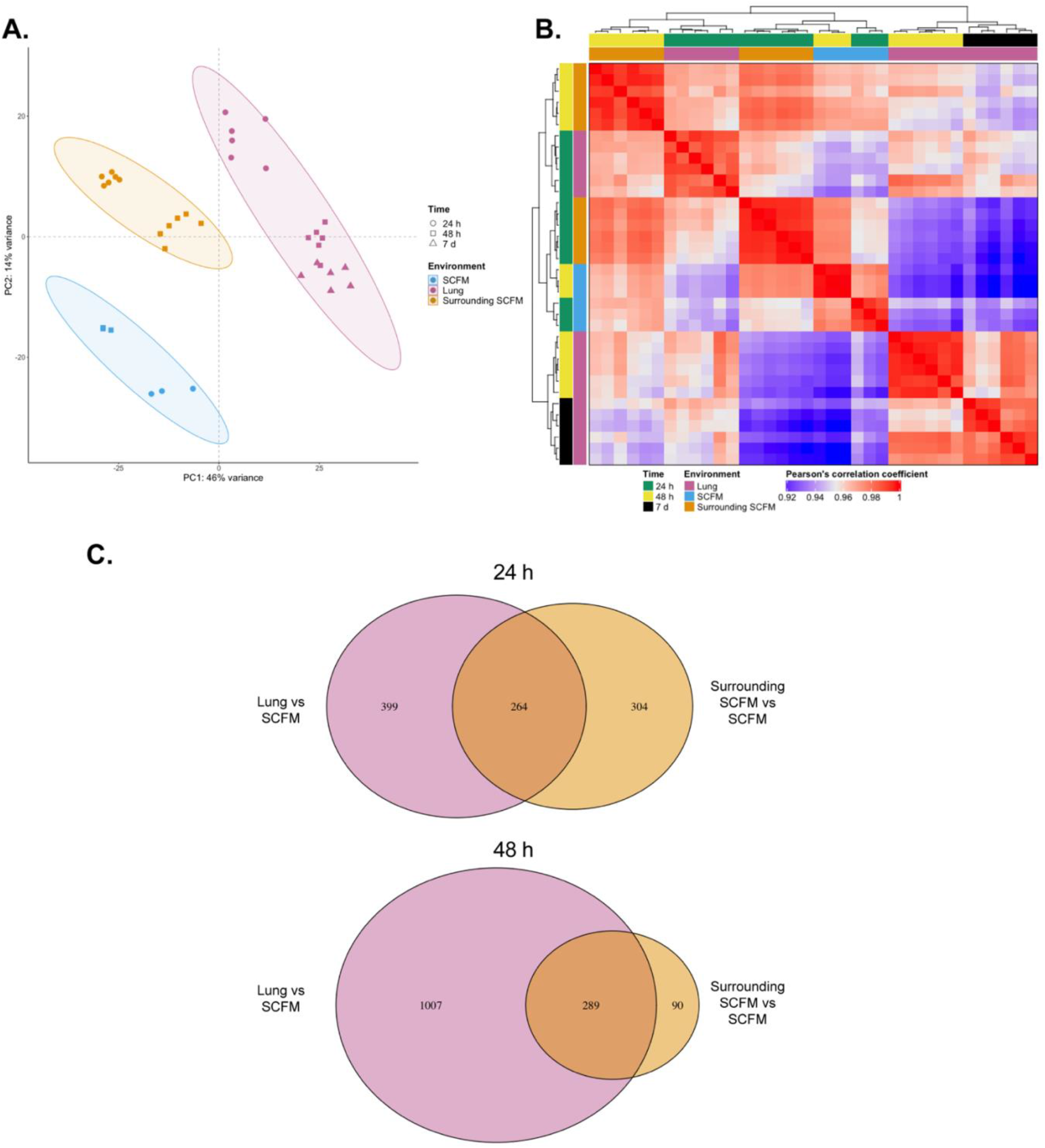
Initial investigation of the *Pseudomonas aeruginosa* PA14 transcriptome across 3 environments: *in vitro* synthetic cystic fibrosis sputum media (SCFM) and the two niches in the *ex vivo* pig lung model: lung tissue surface (lung) and the SCFM surrounding the lung tissue (surrounding SCFM), based on the whole genome (n = 5829 genes). **(A)** Principal component analysis (PCA) considering all genes. Each environment is shown by a different colour and each time point shown by different shaped points (see legend). The 95% confidence ellipses are shown. **(B)** Heatmap showing hierarchical clustering analysis and the Pearson’s correlation coefficient value between each sample (all *r* > 0.9). The *P. aeruginosa* PA14 growth environment and infection time for each sample is shown by different combinations of colours (see legend). **(C)** Venn diagrams of the number of significant differentially expressed *P. aeruginosa* PA14 genes (DEGs) from each contrast, using threshold values of *P* < 0.05 and log_2_ fold change ≥ |1.5|. The shared DEGs are genes that are either underexpressed or overexpressed in both the lung and surrounding SCFM versus *in vitro* SCFM. Genes that were significant DEGs in both contrasts at each time point, but in opposite directions, are not considered to be shared between both contrasts.

Differential expression analysis was then performed to identify significant differentially expressed genes (DEGs) for each contrast (*P* < 0.05, log_2_ fold change ≥ |1.5|). Comparison of the two *ex vivo* environments (lung and surrounding SCFM) and *in vitro* SCFM at 24 h and 48 h PI found a number of significant DEGs shared between lung-associated biofilm and surrounding SCFM. There were also many significant DEGs specific to each EVPL model environment (Fig 1C). Similar numbers of genes were significantly differentially expressed in both the lung-associated *P. aeruginosa* biofilm population and surrounding SCFM, compared with *in vitro*, at 24 h PI (663 and 568 genes respectively). Of these, 264 significant DEGs were common to both populations. At 48 h PI there was a comparable number of DEGs shared between the lung and surrounding SCFM, versus *in vitro* SCFM, to 24 h (Fig 1C). However, the overall number of significant *P. aeruginosa* DEGs in the lung tissue biofilm compared with *in vitro* SCFM had increased (1296 genes) (Fig 1C). This trend was not observed in the surrounding SCFM (decrease to 379 overall DEGs: 90 unique). These findings reveal clear differences in PA14 gene expression between the EVPL model and *in vitro* SCFM and indicate that this difference grows as the biofilm establishes over time on the lung tissue surface. The difference is maintained from 48 h to 7 d PI, with these time points appearing to be more comparable in the number of significant DEGs than 24 h versus 7 d (Fig S5). In contrast, the significant difference between *in vitro* growth and the surrounding SCFM population appears to reduce over time. This indicates that as a chronic *P. aeruginosa* biofilm infection establishes in the EVPL model, the difference between the PA14 transcriptome in the surrounding SCFM and *in vitro* SCFM becomes less distinct, whilst the difference between lung-associated biofilm and *in vitro* becomes more distinct.

### Gene expression differences in EVPL tissue-associated *P. aeruginosa* biofilms versus *in vitro* SCFM growth demonstrate functional importance

Differential expression analysis found significant differences in the *P. aeruginosa* transcriptome depending on whether it was growing as a biofilm associated with pig lung tissue, in the SCFM surrounding the lung tissue or *in vitro* in SCFM. We therefore determined the functional importance of these differences at the two time points that we were able to study across all 3 environments: 24 h and 48 h PI. KEGG pathway enrichment analysis identified multiple pathways that were significantly enriched (*P* < 0.05) in the contrasts between both EVPL environments and *in vitro* SCFM growth at 24 h PI, including quorum sensing (both *P* < 0.05: see Fig S6 for full results). There were also significant KEGG pathways unique to each comparison: cationic antimicrobial peptide resistance and biofilm formation were significantly enriched in the lung-associated biofilm compared with *in vitro* SCFM (both *P* < 0.05). The results indicated that the significant DEGs between comparisons have functional context that may cause different infection phenotypes.

Gene ontology (GO) term analysis was then performed to provide more detailed functional information (significantly enriched: *P* < 0.05). Figure 2 shows the PA14 biological process GO terms that were significantly enriched in the lung-associated biofilm and surrounding SCFM compared with *in vitro* SCFM, at 24 h and 48 h PI. The number of significantly enriched GO terms increases over time in the lung versus *in vitro* SCFM, however it decreases in the surrounding SCFM comparison (Fig 2). This is consistent with our previous conclusion that there was a reduction in number of significant DEGs in surrounding SCFM versus *in vitro* SCFM from 24 h to 48 h PI, and an increase in significant DEGs in the lung-associated biofilm (Fig 1C). Additionally, the GO term ‘response to abiotic stimulus’ was significantly enriched in the surrounding SCFM compared with *in vitro* SCFM 48 h PI (*P* = 0.04, Fig 2D). All associated genes were upregulated in the surrounding SCFM. It is possible this is a result of interaction at the interface between the SCFM and the SCFM-agarose pad placed in the tissue culture plates to support the bronchiolar tissue section. The same 24-well plates were used for *in vitro* growth, so the plate surface is a consistent condition and unlikely to cause expression differences. This GO term was not found to be significantly enriched in the lung-associated PA14 biofilm at either time point (Fig 2A,C).

**Figure 2.**
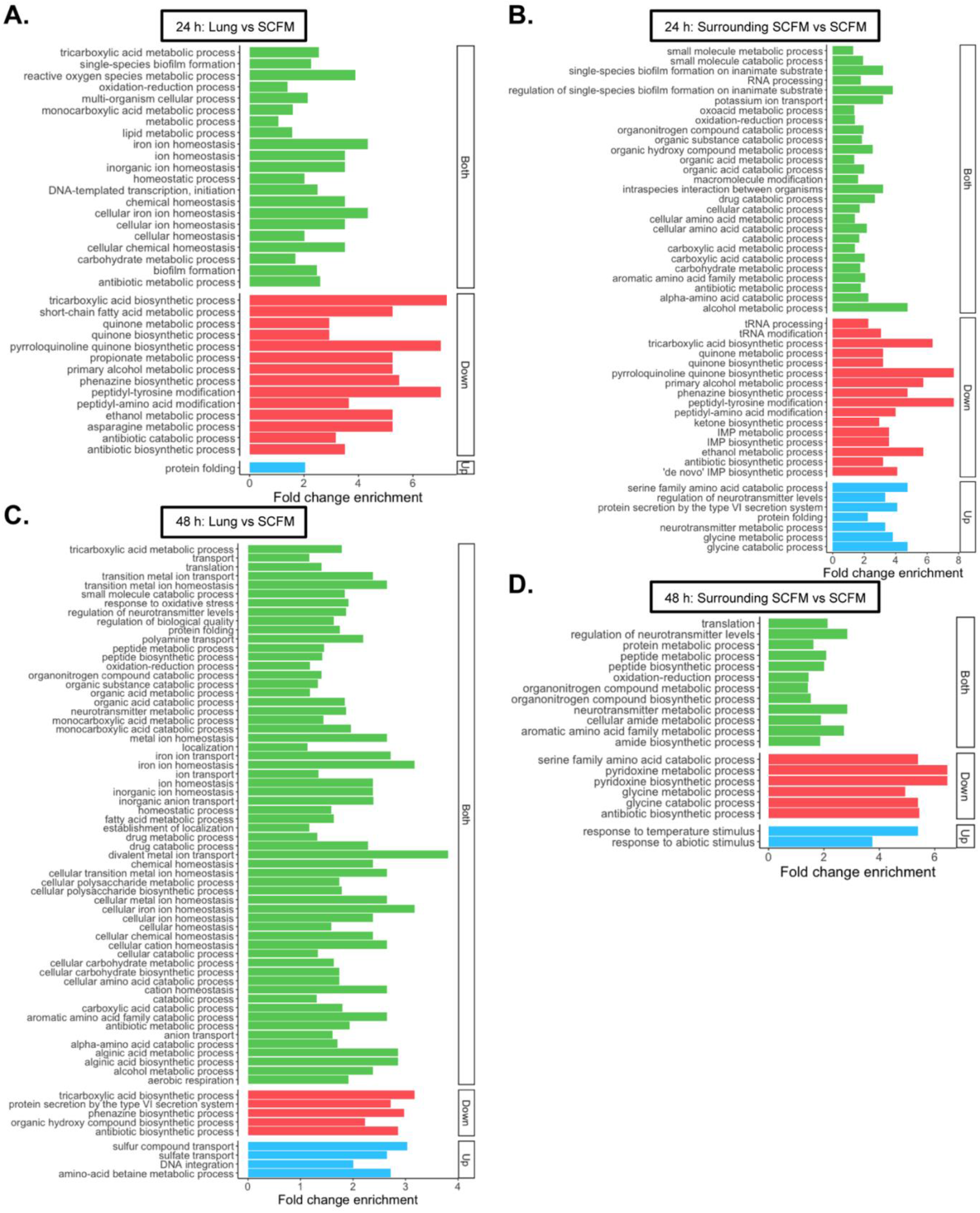
Significantly enriched *Pseudomonas aeruginosa* PA14 biological processes gene ontology (GO) terms (*P* < 0.05, log_2_ fold change ≥ |1.5|) from contrasts between growth from each of the two *ex vivo* pig lung model locations: the lung tissue (lung) and the synthetic cystic fibrosis sputum media (SCFM) surrounding the lung tissue (surrounding SCFM), and *in vitro* SCFM (SCFM). The analysis was performed on samples from 24 h post infection (**A, B**) and 48 h post infection (**C, D**). Each graph shows the significantly enriched biological processes GO terms for the particular contrast on the y-axis and fold change enrichment on the x-axis. The fold change enrichment is the fold difference in expression of significant differentially expressed genes (DEGs) in the analysis associated with that GO term than expected by random chance. Green bars show terms where there were some associated significant DEGs overexpressed and some underexpressed (both). The red bars show terms where all associated significant DEGs in the contrast were underexpressed and blue bars show terms where all associated significant DEGs were overexpressed.

The GO term ‘phenazine biosynthetic process’ was found to be significantly enriched in the *P. aeruginosa* PA14 lung biofilm versus *in vitro* SCFM growth at both time points (Fig 2A,C). The KEGG pathway ‘phenazine biosynthesis’ was also found to be significantly enriched for these comparisons (Fig S6). All genes for the biosynthesis of the exotoxin pyocyanin were downregulated in the lung-associated biofilm at 48 h compared with *in vitro* SCFM (Fig S6). This suggests that phenazine biosynthesis, particularly pyocyanin production, may be an important differentiation between *in vitro* and pig lung grown *P. aeruginosa* biofilms.

### *P. aeruginosa* quorum sensing-associated pathways are downregulated in the EVPL model compared with *in vitro* SCFM growth, as observed in CF sputum samples

Quorum sensing (QS) pathways in *P. aeruginosa* have long been considered to play a role in the establishment of biofilms [18]. Three key *P. aeruginosa* QS systems are associated with controlling biofilm formation in human infection: the LasI/R system, RhlI/R and the *Pseudomonas* quinolone signal (PQS) [12,19,20]. These acyl-homoserine lactone (AHL) associated systems regulate the expression of numerous genes associated with virulence and chronic biofilm infection [12]. We found that the QS KEGG pathway was significantly enriched in the SCFM surrounding *P. aeruginosa* infected EVPL tissue versus *in vitro* SCFM growth at 24 h and 48 h PI, and in the lung biofilm versus *in vitro* at 24 h PI (Fig S6). The majority of the significant DEGs in this pathway were under-expressed in the lung environments compared with *in vitro* (Table 1). These results, alongside QS being an extensively researched area with the potential as a new therapeutic target in CF [21,22], led to more detailed exploration of the expression of QS-associated genes in the EVPL model.

Further investigation of the QS-associated significant DEGs demonstrated that most of these genes are in the phenazine biosynthesis operons, as shown in Table 1 (for full list see Table S3). This finding further supports these pathways as key to the difference between *ex vivo* and *in vitro* grown *P. aeruginosa* transcriptomes. These genes had a greater log_2_ fold change reduction on the lung-associated biofilm compared with *in vitro* SCFM at 24 h and 48 h PI, than surrounding SCFM versus *in vitro*. The expression of *pqsC* and *pqsD*, both important in the production of PQS [23], was also found to be underexpressed in the lung compared with *in vitro* SCFM 48 h PI (Table 1). This suggests that this third QS system is also being downregulated over time in the lung-associated biofilm in comparison with *in vitro* SCFM.

The study of *P. aeruginosa* transcriptomes by Cornforth *et al.* (2018) found that expression of a set of conserved genes under las-regulation differs between *in vitro* growth and human infection, with lower levels of expression observed *in vivo* [8]. This set of 42 *P. aeruginosa* genes regulated by the las QS system were found to be conserved amongst CF lung isolates [25]. Thus, these genes were specifically investigated in our RNA-seq results. The expression of the equivalent gene set in *P. aeruginosa* PA14 when grown on *ex vivo* pig lung tissue compared with *in vitro* SCFM showed similar significant differential expression to CF sputum versus *in vitro* as found by Cornforth *et al*.[8] (Fig 3). At 24 h PI similar changes in expression of these genes were found in the EVPL tissue-associated biofilm compared with SCFM *in vitro* (Fig 3A), as seen in human CF sputum (*lasA*: −2.09 & −3.38 log_2_ fold change respectively and *lasB*: −3.35 & −3.30). At 48 h PI, the expression profile of this gene set becomes even more comparable to the relative expression in CF sputum (Fig 3B). These genes were also studied in surrounding SCFM growth versus *in vitro* SCFM at both time points and although the pattern was less obvious, there were some similarities with expression in CF sputum versus *in vitro* (Fig S7). These findings support the conclusion that both environments of the EVPL model more closely capture a key aspect of human CF *P. aeruginosa* infection than *in vitro* growth, but the lung-tissue associated biofilm at 48 h appears to be most representative.

**Figure 3.**
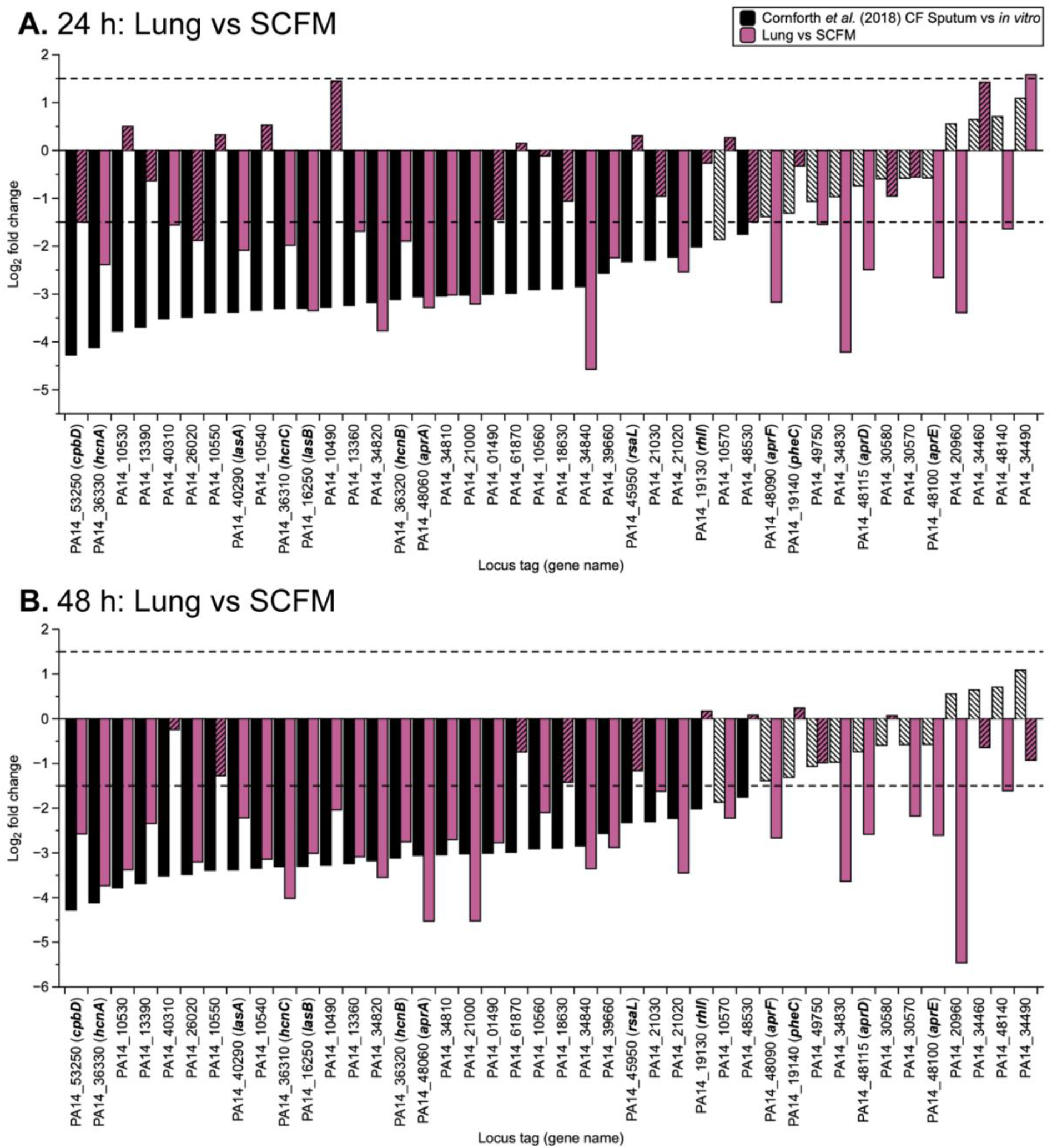
The log_2_ fold change (lfc) in expression of 42 *Pseudomonas aeruginosa* quorum sensing genes, controlled by the las regulon, conserved in human infection. The expression of *P. aeruginosa* PA14 grown on the lung tissue of the *ex vivo* pig lung tissue (Lung) versus *in vitro* synthetic cystic fibrosis media (SCFM) at 24 h (**A**) and 48 h (**B**) post infection are shown by the purple bars. Each graph also includes expression of the gene set by *P. aeruginosa* from human cystic fibrosis sputum versus *in vitro* conditions taken from Cornforth *et al.* [8], shown by the black bars. The locus tags shown are for *P. aeruginosa* PA14 with gene names in bold where appropriate. Bars with the striped fill are not significantly differentially expressed for that contrast (*P* < 0.05, lfc ≥ |1.5|). The dashed lines represent the threshold lfc value for differential expression.

Alongside our RNA-seq findings, we performed phenotypic assays to measure the concentration of extracellular 3-oxo-dodecanoyl homoserine lactone (3-oxo-C12-HSL) in each environment at 24 h and 48 h PI. Production of 3-oxo-C12-HSL is encoded by the gene *LasI*, it binds to the LasR transcriptional regulator to regulate expression of numerous genes [20]. We found a significantly lower concentration of extracellular 3-oxo-C12-HSL in both *ex vivo* conditions compared with *in vitro* SCFM at 48 h PI (Fig S8), consistent with the downregulation of the *las*-controlled gene set. However, neither *lasI* nor *lasR* were found to be significantly differentially expressed for any comparisons (Table S3). A previous study by Aendekerk *et al.* [26] that investigated the MexGHI-OpmD efflux pump found that mutations in this pump resulted in *P. aeruginosa* being unable to produce 3-oxo-C12-HSL. RT-PCR showed that this lack of 3-oxo-C12-HSL production did not correlate with inhibition of *lasI* or *rhlI* transcription, thus MexGHI-OmpD exerts its effects on 3-oxo-C12-HSL levels post-transcriptionally. This mechanism is consistent with our results, as we found that all genes encoding MexGHI-OpmD were significantly underexpressed in the lung-associated biofilm compared with *in vitro* SCFM growth at 48 h PI (Fig 4A). Genes associated with self-degradation of acyl homoserine lactone (AHL) signals were not found to be significantly over expressed in any of our comparisons (Table S4). The expression of the MexAB-OprM pump components, which transports 3-oxo-C12-HSL out to the cell, were also not significantly different in the lung associated-biofilm (Fig 4C), supporting the conclusion that the difference in extracellular 3-oxo-C12-HSL concentration is a result of reduced production associated with MexGHI-OpmD activity.

**Figure 4.**
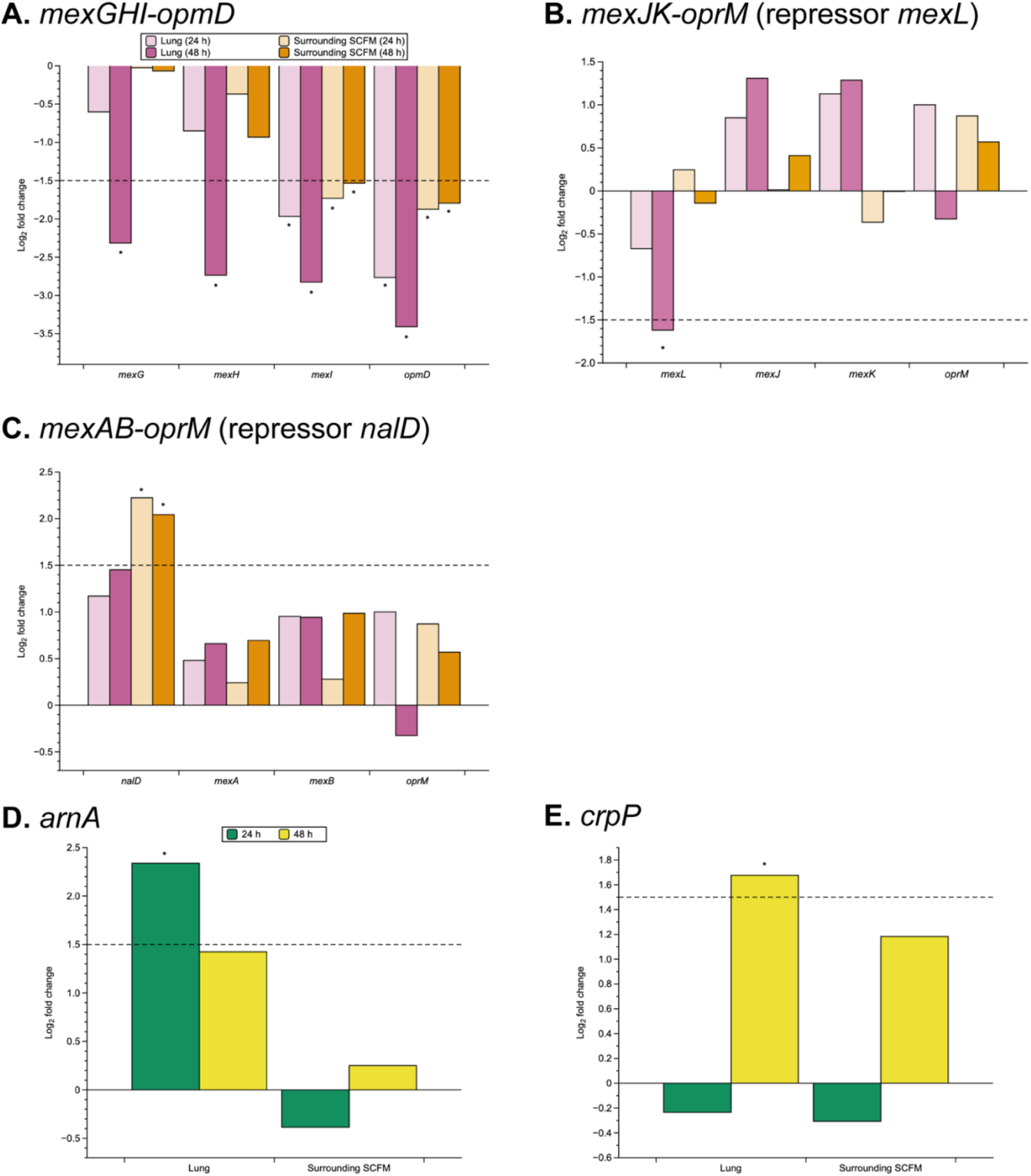
The log_2_ fold change (lfc) of genes of interest associated antibiotic resistance predicted by the Comprehensive Antibiotic Resistance Database (CARD) [27]. Comparisons of gene expression were performed for the two environments of the *ex vivo* pig lung model: the **lung** tissue and **surrounding** synthetic cystic fibrosis media (**SCFM**) versus *in vitro* **SCFM** at two timepoints (24 h and 48 h post infection). The dashed lines represent the threshold lfc value for a gene to be considered significantly differentially expressed (lfc ≥ |1.5|), and for comparisons where this was statistically significant (*P* < 0.05) are denoted with a *. Each bar colour represents a different comparison and time point (see legends). **(A-C)** The lfc values for each comparison of efflux pumps and any repressors where significant expression differences were found in at least one comparison. **(D-E)** The lfc values for each comparison of individual genes where significant expression differences were found in at least one comparison.

### Genes associated with antibiotic resistance are differentially expressed in EVPL environments compared with *in vitro* SCFM

Following investigation of quorum sensing gene expression and the implication on efflux pump expression, we aimed to determine whether growth environment alone – with no prior exposure to antimicrobials – resulted in differing expression of antimicrobial resistance (AMR) associated genes. We have previously shown that high levels of antibiotic tolerance are evident in the EVPL model [16] and clinical infection typically demonstrates much higher resistance to antimicrobial treatment than is predicted *in vitro*. We looked at the expression of 52 *P. aeruginosa* PA14 AMR genes, predicted using the Comprehensive Antibiotic Resistance Database (CARD) [27], in each contrast. There was significant differential expression of 22 AMR associated genes in the two EVPL model environments: lung tissue and surrounding SCFM, compared with *in vitro* SCFM growth of PA14 (see Table S5).

Three efflux pumps were of particular interest in at least one of the comparisons (Fig 4A-C). All genes encoding the efflux pump MexG/HI-OpmD were found to be significantly underexpressed in the lung-associated biofilm compared with *in vitro* SCFM growth at 48 h PI (Fig 4A). MexHI-OpmD is associated with resistance to fluoroquinolones, whereas MexGHI-OpmD is associated with the post-transcriptional regulation of 3-oxo-C12-HSL production as described above, and exerts effects on antibiotic sensitivity via its effects on QS [28]. The transcriptional repressor of the MexJK-OprM efflux pump (*mexL*), linked to triclosan resistance [29], was also found to be significantly underexpressed in the lung-associated biofilm compared with *in vitro* SCFM (Fig 4B). Conversely, a transcriptional repressor of the MexAB-OprM efflux pump (*nalD*), was significantly overexpressed in the surrounding SCFM at 24 h and 48 h compared with *in vitro* SCFM. Previously, mutations in repressors of this efflux pump have been linked to carbapenem-resistance and over-expression of MexAB-OprM [30]. However, this pump has also been proven to no longer be required upon the formation of a mature biofilm [31], which may suggest why a transcriptional repressor is being overexpressed in the surrounding SCFM as a *P. aeruginosa* biofilm is establishing in the EVPL model [15].

Two individual genes of interest were found to be significantly differentially expressed in at least one comparison (Fig 4D-E). Both genes (*arnA* and *crpP*) are associated with the *arn* locus, linked with resistance to cationic antimicrobial peptides (CAMPs) [32]. They were both found to be significantly differentially expressed in the lung tissue-associated biofilm compared with *in vitro* SCFM: *arnA* overexpressed at 24 h PI and *crpP* at 48 h. This was consistent with minimum inhibitory concentration (MIC) values for colistin and polymyxin B (CAMPs) of *P. aeruginosa* PA14 cells from the lung tissue-associated biofilm found to be increased compared with SCFM and the standard CAMHB at both time points (Table S6). These findings indicated that growth of PA14 biofilm on EVPL tissue causes changes in expression of AMR pathways compared with *in vitro* SCFM that have a phenotypic effect.

## Discussion

*P. aeruginosa* infection of the CF lung is one of the most well-studied biofilm infection contexts, yet it remains highly resistant to the most aggressive antimicrobial treatments available, and eventually results in loss of lung function and accelerated death [1,3]. Research to improve prevention measures and treatment of chronic CF infection is hindered by the lack of accurate laboratory growth conditions and models to truly mimic biofilm formation as observed in human infection. It has been shown that growth environment plays an integral role in the *P. aeruginosa* transcriptome; *P. aeruginosa* gene expression in CF sputum samples has a distinct transcriptional profile compared with current laboratory models including mice models and an enhanced, updated version of SCFM (SCFM2) [8–10]. We compared gene expression in our EVPL model, considering the tissue-associated biofilm and bacteria in the surrounding SCFM as separate populations, with gene expression in SCFM *in vitro*. We focused on key aspects of infection that may not be well captured by current laboratory models: quorum sensing and antibiotic resistance. We propose that our previously described EVPL model can be used to mimic *P. aeruginosa* biofilm infection as seen in CF not only in structure [15], but also in key aspects of the transcriptome.

We compared gene expression across the whole genome in both environments of the EVPL model, and in SCFM *in vitro,* across multiple time points. Our analyses revealed that the lung tissue and surrounding SCFM of the EVPL model create two distinct niches from each other and from *in vitro* cultures. To date, development of laboratory model systems has predominantly focused on recapitulating CF sputum through production of artificial sputum media [33]. However, sputum media such as SCFM and SCFM2, discussed above, do not attempt to recapitulate the tissue-associated biofilm infection that also occurs in the CF airway. Our results show a clear difference in *P. aeruginosa* gene expression between a synthetic sputum and lung tissue of the EVPL model. This highlights the tissue-associated biofilm as a distinct niche that is arguably just as important to further understand infection dynamics as the ‘sputum-like’ population. A clear distinction in gene expression changes in the population over time in *ex vivo* conditions versus *in vitro* was also found. Thus, as well as growth environment being important for expression differences at each point of comparison, it also affected the dynamics of infection progression.

The greatest difference between *in vitro* and *ex vivo* conditions was in the lung tissue-associated biofilm (versus *in vitro* SCFM) at 48 h PI. In particular, genes in phenazine biosynthetic pathways were found to differ in expression. Phenazines are redox-active pigments with antimicrobial activity that are known to cause changes in gene expression and antibiotic susceptibility [34]. The phenazine pyocyanin is an important *P. aeruginosa* virulence factor and all genes in its biosynthetic pathway were found to be under-expressed in the lung tissue environment at 48 h compared with *in vitro*. Virulence factor production is typically downregulated during the chronic stages of infection, in fact overproduction of pyocyanin during CF infection is considered an unusual phenotype [35]. Thus, these findings indicate that the EVPL model is driving *P. aeruginosa* into a chronic state over time that is not seen *in vitro*. This may allow for study of chronic, established biofilms, which are the typically difficult to treat stage of infection.

Phenazines are also signalling molecules and regulation of the associated operons involves all three *P. aeruginosa* QS systems: *las*, *rhl* and *pqs* [36]. QS has been shown to be a key aspect of infection where current laboratory studies cue overexpression upon comparison to human infection [8]. We found that the shift in expression of conserved *las* controlled genes in the lung-associated biofilm versus *in vitro* SCFM at 48 h PI is almost identical to the shift in expression of these genes in CF sputum versus *in vitro* [8]. These results suggest that the EVPL model could more accurately mimic the QS expression profile seen in human infection than other current laboratory models. While QS is important in the initial stages of infection, an accumulation of mutations in QS pathways are observed over time in CF, suggesting that it is not as active in the later stages of infection [11]. Interestingly, *lasI/R* and *rhlI/R* were not found to be significantly differentially expressed in any of our comparisons – the differences were in downstream las- and rhl-regulated genes. A reduced concentration of extracellular 3-oxo-C12-HSL produced by *P. aeruginosa* was also measured in the *ex vivo* environments compared with *in vitro* SCFM growth at 48 h PI. *P. aeruginosa* mutants for the efflux pump-associated genes *mexI* and *opmD* have previously been shown to not produce 3-oxo-C12-HSL due to an intracellular accumulation of a toxic PQS precursor due to loss of pump activity [26]. We found that both genes were downregulated in the lung and surrounding SCFM versus *in vitro* SCFM growth, indicating that the reduction in production of the QS molecule is likely caused by reduced expression of the efflux pump. Interestingly, genes that encode amidases able to degrade AHL signals [37] (PA14 homologues of *pvdQ*, *quiP*, *hacB*) were not found to be differentially expressed between the EVPL model and SCFM at either time point. Thus, our results indicate that the reduction in extracellular 3-oxo-C12-HSL is not due to quorum quenching activity and is likely a reduction in production driven by the downregulation of MexGHI-OpmD. Hence, the differences in QS expression between *P. aeruginosa* grown in the EVPL model and *in vitro* SCFM may be part of a wider network that also has implications for antibiotic resistance.

Increased antibiotic resistance is a characteristic of *P. aeruginosa* infection of the CF lung that poses significant clinical concern. We found a number of efflux pump genes and resistance-associated operons to be significantly differentially expressed in the EVPL model. The downregulation of the MexGHI-OpmD efflux pump in the tissue-associated biofilm at 48 h, as discussed above, may be linked to increased antibiotic resistance as well as QS effects. Mutations in *mexI* and *opmD* have previously been shown to increase *P. aeruginosa* resistance to aminoglycosides, β-lactams and quinolones [26]. A transcriptional regulator of the MexAB-OprM pump (*nalD*) was found to be overexpressed in the surrounding SCFM compared with *in vitro* SCFM, indicating a reduction in this pump also. However, no significant difference in any of the efflux pump genes or the repressor were found in the lung-associated biofilm, as was the case for the other efflux pumps found. The spatial arrangement of MexAB-OprM has been previously proven to be heterogenous within infection populations, with a higher proportion of the pump found in cells that formed a dense biofilm [31]. This may explain why our results indicate there are higher levels of the efflux pump in the lung-associated population than surrounding SCFM. This further distinction between the surrounding SCFM and lung tissue suggests these two spatially linked environments promote different patterns of gene expression that may result in varying antibiotic resistance phenotypes. The model provides a heterogenous infection population that may express different mechanisms of resistance that cannot be captured by *in vitro* SCFM alone. This is an important consideration for determining treatment approaches and novel treatments for *P. aeruginosa* infection in the CF lung.

The over-expression of *arnA* in the lung-associated biofilm at 24 h PI, and *crpP* at 48 h is also of clinical interest, as both genes are linked to CAMP resistance. The gene *arnA* is part of the arn locus that functions via lipid A modifications increasing resistance [32]. Upregulation of this gene at 24 h PI but not 48 h may be providing resistance prior to biofilm formation, after which point the biofilm matrix acts as a protection mechanism against antibiotic treatment. In contrast, *crpP* is upregulated at 48 h PI. CrpP has been shown to confer resistance to ciprofloxacin via phosphorylation [38]. The upregulation of this gene in the tissue-associated biofilm despite no antibiotic treatment suggests this pathway is initiated by growth environment not antibiotic exposure. As well as providing further insight into resistant infection, these findings highlight the importance of using a clinically relevant infection model, such as the EVPL, to better understand the resistance profile of bacteria *in vivo*. Conversely, we must consider that people with CF receive numerous antibiotic treatment courses, so the resistance profiles of clinical infection will be impacted by this as well as the host environment.

In conclusion, we have demonstrated that the EVPL model creates two environments of interest: tissue-associated biofilm and the surrounding SCFM, that are distinct from *in vitro* SCFM. Gene expression in the tissue-associated biofilm at 48 h appears to be more representative of an established, chronic infection of the CF lung. We have focused on expression at a whole population level, however future expression studies at the single-cell level may reveal further insight into the infection dynamics and population heterogeneity to improve development of novel, effective treatments. Future studies could also focus on transcriptomics of clinical *P. aeruginosa* isolates in the EVPL model, as their response to the environment may differ from that of PA14 [9]. These studies have the capacity to be conducted over longer timescales than studies conducted on *P. aeruginosa in vitro* in SCFM; by 7 days PI, we could not recover mRNA from cells grown in vitro and bacteria appeared stressed, but EVPL-grown cells were viable and yielded good quality mRNA. Overall, the EVPL model is a useful platform to dissect key aspects of *P. aeruginosa* pathophysiology in CF that is not possible in current *in vitro* models.

## Materials and Methods

### Bacterial strain

The wild type (WT) *Pseudomonas aeruginosa* PA14 strain used in this study was from the University of Washington [39,40]. It was grown on Luria-Bertani (LB) agar (Melford Laboratories) for 24 h at 37 °C prior to all infections and to determine colony forming units (CFU). The pSB1075 *Escherichia coli* biosensor, which carries a fusion of *lasRI*’::*luxCDABE* [41] was used to measure 3-oxo-C12-HSL in culture supernatants.

### Synthetic cystic fibrosis sputum medium

The synthetic cystic fibrosis sputum medium (SCFM) was prepared based on a previously published recipe [33], with the glucose removed. Preliminary work found that with the addition of porcine lung tissue, glucose facilitated endogenous bacterial growth, and the presence or absence of glucose did not affect *P. aeruginosa* growth [14].

### *Ex vivo* pig lung dissection and infection

All pig lungs used were supplied by a commercial butcher (Steve Quigley & Sons, Cubbington, UK) and dissected on the day of arrival from the abattoir. *Ex vivo* pig lung (EVPL) tissue was dissected to extract the bronchioles as previously described [14,15,42]. Following ultraviolet (UV) light sterilisation, the square bronchiolar tissue pieces were placed into each well of a 24-well plate/s with a 400 µl, UV-sterilised, 0.8% (w/v) agarose pad.

To infect each tissue piece, a 29 G sterile hypodermic needle (Becton Dickinson Medical) was used to touch the surface of a colony of the infection strain from an overnight LB plate and then transferred to the tissue by lightly piercing the surface. To ‘mock inoculate’ the uninfected negative control pieces a sterile needle was used. To fully replicate the CF lung environment, 500 µl SCFM was added to each tissue-containing well and the plate covered with a Breathe-Easier® membrane (Diversified Biotech) sterilised with UV. Plates were incubated stationary for the required length of time at 37 °C.

### Bacterial recovery from the EVPL model and bacterial count determination

There were two environments in the EVPL model: the bronchiolar tissue (tissue) and the SCFM surrounding each tissue piece (surrounding SCFM). To recover bacteria from the tissue, each tissue piece was removed from the 24-well plate following incubation and transiently washed in 500 µl phosphate-buffered saline (PBS). Tissue sections were then placed in sterile homogenisation tubes (Fisherbrand), each had eighteen 2.38 mm metal beads (Fisherbrand) and 1 ml PBS added. To recover the biofilm-associated population, the tissue-containing tubes were bead beat using a FastPrep-24 5G (MP Biomedicals) for 40 s at 4 m s^−1^. The surrounding SCFM was directly transferred to individual sterile 2 ml DNA lo-bind tubes from the 24-well plate (Eppendorf).

To determine bacterial load, an aliquot was taken from each tissue homogenate and surrounding SCFM sample and serially diluted in PBS. Dilutions were plated on LB agar and incubated at 37 °C for 24 h. Tissue and surrounding SCFM taken from the same sample were recorded as the same repeat number for comparison. Colony counts were performed and the CFU per lung and per ml for the surrounding SCFM were calculated.

### Microbial cell viability from EVPL tissue

The BacTiter-Glo™ microbial cell viability assay (Promega) was used on EVPL tissue-associated samples across a 7 d infection period. Cell viability was measured by the amount of ATP (nM) produced. Following lung dissection, infection, incubation and recovery, the lung homogenate was equilibrated to room temperature. Once room temperature was reached, 100 µl of each sample was added to each well of a 96-well black plate. The assay was performed as per kit instructions using a Tecan Spark 10M multimode plate reader and ATP (nM) was determined using the ATP (Jena Bioscience) standard curve produced.

### P. aeruginosa in vitro SCFM growth

*P. aeruginosa* colonies from an overnight LB agar plate were suspended in an appropriate volume of SCFM to an OD_600nm_ of 0.05. Aliquots of 1 ml were added to each well of a 24-well plate and covered with a UV-sterilised Breathe-Easier® membrane. Plates were incubated stationary at 37 °C for the required length of time.

### Production of 3-oxo-C12-HSL

The lung homogenate, surrounding SCFM and *in vitro* SCFM cultures were filter sterilised using a 0.2 µm pore syringe filter (Fisherbrand™) into sterile 2 ml Eppendorf tubes. The sterile supernatants were then stored at −20 °C prior to performing the assay.

The *E. coli* pSB1075 bioreporter for 3-oxo-C12-HSL was grown in 10 ml LB broth (+ 10 µg ml^−1^ tetracycline) overnight at 37 °C, with 170 rpm shaking. The overnight culture was then diluted 1:100 in 15 ml LB broth and incubated at 37 °C, with 170 rpm shaking for 3.5 h. The culture was centrifuged at 13,000 rpm for 2 min and the supernatant discarded. The pellet was then re-suspended in 15 ml PBS. The centrifugation and re-suspension were repeated twice more, and the final pellet resuspended in 15 ml LB. The OD_600nm_ was adjusted to 0.1 using LB broth. Sterile sample supernatants were defrosted on ice, diluted 1 in 10 in PBS, and mixed 1:1 with the final *E. coli* culture to a volume of 200 µl, in a 96-well black plate. A standard curve was also performed in the same plate, using 3-oxo-C12-HSL (Sigma-Aldrich) concentrations from 1 nM to 0.000001 nM in ten-fold increments. The plate was then incubated at 37 °C for 7.5 h in a Tecan Sark 10M multimode plate reader and the OD_600nm_ and relative light units (RLU) were read every 15 min. The RLU/OD was calculated, and the final concentrations of 3-oxo-C12-HSL were determined at the inflection point, using the standard curve values.

### Minimum inhibitory concentration assay

Lung homogenate was diluted 1 in 100 in SCFM to be used as the bacterial inoculum to perform minimum inhibitory concentration assays (MICs). The inoculum for MICs in SCFM and cation-adjusted Mueller Hinton broth (CAMHB) were prepared as a MacFarland standard. Briefly, *P. aeruginosa* PA14 was grown on an LB agar plate overnight and colonies were suspended in the relevant media to an OD_600nm_ of 0.08 – 0.1, and then diluted 1 in 100.

MICs were performed following the broth microdilution method as described by Wiegand *et al*. [43]. The antibiotics, colistin (Acros Organics) and polymyxin B (Sigma-Aldrich), were serially diluted 1 in 2, from 128 – 0.0125 µg ml^−1^, to a final volume of 100 µ, in tissue culture (TC)-treated 96-well plates (Corning) using the relevant media. Subsequently, 100 µl of bacterial inoculum was added to each antibiotic well as well as positive control wells with no antibiotic added. Negative control wells were prepared with only water. Plates were then incubated at 37 °C for 18 h. The lowest antibiotic concentration where no growth was visible was recorded as the MIC value.

### *P. aeruginosa* RNA extraction

*P. aeruginosa* RNA was extracted from three infection environments: EVPL tissue, EVPL surrounding SCFM and *in vitro* SCFM. Each culture of interest was transferred to individual sterile 2 ml DNA lo-bind tubes: 1 ml *in vitro* culture per tube, 500 µl surrounding SCFM per tube and 1 ml lung homogenate per tube. Subsequently, 0.5 volume sterile killing buffer (20 mM Tris-HCl pH 7.5, 5 mM MgCl_2_, 20 mM NaN_3_) was added to each tube and then centrifuged at 13,000 rpm for 1 min. The samples were snap frozen and stored at −80 °C for at least 1 h.

Samples were defrosted on ice and the supernatant gently removed, ensuring the pellet remained in-tact. Each pellet was resuspended in 600 µl sterile LETS buffer (0.1 M LiCl, 0.01 M Na_2_EDTA, 0.01 M Tris-HCl pH 7.5, 0.2% SDS) and transferred to 2 ml lysing matrix B tubes (MP Biomedicals). Tubes were bead beat using a FastPrep-24 5G for three cycles: 6 m s^−1^ for 40 s then 5 min incubation on ice. The samples were further incubated in the lysing matrix B tubes until they reached room temperature, centrifuged for 10 min at 13,000 rpm and 600 µl 125:24:1 Phenol Chloroform Isoamyl alcohol (PCI) pH 4.5 (Invitrogen) was added. Each tube was vortexed for 5 min at ∼14,000 rpm then centrifuged at 15,000 rpm for 5 min at 4 °C. The top layer of solution was transferred to sterile 2 ml RNase-free tubes (Sarstedt Ltd) and 1 volume of 125:24:1 PCI pH 4.5 added. The previous vortex, centrifuge and top layer transfer steps were repeated and 1 volume 24:1 Chloroform Isoamyl alcohol (CI) (Sigma-Aldrich) was added. All samples were then centrifuged at 15,000 rpm for 5 min and the top layer transferred to a new 2 ml RNase-free tube. Finally, 0.1 volume of 3M NaCH_3_COO pH 5.2 and 1 volume of isopropanol was added and mixed by inverting each tube. Samples were stored at −20 °C overnight.

### RNA precipitation and DNA removal

RNA samples were defrosted on ice and centrifuged at 15,000 rpm for 15 min. The supernatant was removed, and each pellet resuspended in 1 volume 70% (v/v) ethanol. Resuspended samples were centrifuged at 15,000 rpm for 15 min at 4 °C. Supernatant was removed and pellets left to dry for ∼15 min by a flame, then resuspended in 50 µl RNase-free water. Each sample was incubated on ice for 3 h then for a further 30 min at room temperature. The precipitated RNA concentration was determined using the Qubit™ RNA BR (broad range) Assay kit. All RNA was snap frozen and stored at − 80 °C.

Frozen RNA was defrosted and any samples above 200 µg ml^−1^ were diluted in RNase-free water to ≤ 200 µg ml^−1^. Each sample was transferred into PCR tubes, to a maximum of 50 µl per tube, samples with a higher volume were divided into multiple tubes. A 9:1 ratio of DNase I buffer (10X) (Invitrogen) to RNA sample respectively was added to each tube. Subsequently, 2 µl DNase I (Invitrogen) was added to each reaction and the tubes incubated at 37 °C for 30 min. A further 2 µl DNase I was added, and the incubation repeated. An equal volume ethanol to sample was added, thoroughly mixed then transferred into Zymo-Spin™ ICC columns (Direct-zol™ RNA MiniPrep Plus kit, Zymo Research). Samples separated due to high concentration were combined in one spin column. Spin columns were centrifuged at 13,000 rpm for 30 s and the flow-through discarded. Each RNA sample was cleaned up using the Direct-zol™ RNA MiniPrep Plus kit. For the final elution, tubes were incubated at 55 °C for 5 min to increase RNA yield. A 16s PCR and gel electrophoresis were used to confirm complete DNA removal and for any samples with detectable DNA, the protocol was repeated. Final RNA concentration was confirmed using the Qubit™ RNA BR Assay kit. Samples were snap frozen and stored at −80 °C.

### Quality check of RNA and sequencing

Total extracted RNA quality was confirmed for sequencing using the Agilent 2100 Bioanalyzer system using the RNA 6000 Pico Kit (Agilent). Bacterial and mammalian rRNA depletion, and Illumina library preparation for strand-specific RNA-sequencing was performed by Genewiz then samples were sequenced on an Illumina NovaSeq. 150 bp paired-end run.

### Bioinformatic analyses

RNA-seq reads were initially quality checked using FastQC v0.11.8 [44], which was subsequently used after each step of initial data preparation. Reads were trimmed using Trimmomatic v0.38 [45], with a minimum read length threshold of 25 bp [8]. Any residual bacterial rRNA transcripts, and eukaryotic where appropriate, were filtered out using SortmeRNA v2.1b [46]. All reads from EVPL-grown samples (surrounding SCFM and lung) were mapped to the pig genome (*Sus scrofa*: NCBI, GCF_000003025.6) using HISAT2 v2.1.0 [47] and the aligned reads were removed from the trimmed reads using Seqtk v1.3-r106 [48]. The remaining EVPL sample reads and *in vitro* SCFM sample reads were aligned to the *P. aeruginosa* UCBPP-PA14 genome (NCBI, assembly GCF_000014625.1) using the BWA v0.7.17-r1188 aligner with the MEM algorithm [49]. Reads mapped to coding sequences were counted using the “featureCounts” function from the R package Rsubread v2.0.1 [50] against the *P. aeruginosa* UCBPP-PA14 annotation sourced from *Pseudomonas.com* [24]. Count data was normalised using the rlog Transformation function from the R package DESeq2 v1.26.0 [51], then used for principal component analysis (PCA) based on all genes in the analysis (5829) using the “plotPCA” function within DESeq2. The 95% confidence ellipses were determined and added to the PCA plot using ggpubr v0.4.0 [52]. A hierarchical clustering heatmap was produced to show pairwise correlation values based on Pearson’s correlation coefficient for all sample comparisons using the “HeatmapAnnotation” function from the ComplexHeatmap v2.2.0 R package [53].

Differential gene expression analysis between samples from different growth environments and time points was performed using DESeq2; genes were considered significantly differentially expressed with an adjusted *P*-value < 0.05 (Benjamini-Hochberg procedure to control the false discovery rate) and log_2_ Fold Change (FC) ≥ |1.5|. KEGG pathway enrichment analysis of the differentially expressed genes (DEGs) was carried out with “enrichKEGG” function from the R package clusterProfiler v3.14.3 [54] with the *P. aeruginosa* UCBPP-PA14 organism KEGG code (“pau”). KEGG pathways were considered significantly enriched using an adjusted *P*-value < 0.05 (Benjamini-Hochberg). Gene ontology (GO) term enrichment analysis was also performed using topGO v2.38.1 [55]. Reported significantly enriched GO terms were based on Fisher’s exact test *P-*value < 0.05. Antimicrobial resistance genes investigated were sourced from *Pseudomonas.com* [24], based on predictions by the Comprehensive Antibiotic Resistance Database (CARD) [27].

## Acknowledgments

We would like to thank all at Steve Quigley & Sons butchers for donating all pig lungs used; Leo Eberl, Steve Diggle and Roman Popat for bacterial strains; Jenna Lam for her help with the RNA extraction protocols; Blessing Anonye for her guidance; Freya Allen for performing some of the MIC assays; and Marvin Whiteley, Dan Cornforth and Zamin Iqbal for insightful discussion. We would also like to acknowledge the help of the Media Preparation Facility in the School of Life Sciences at the University of Warwick, with special thanks to Cerith Harries and Caroline Stewart. This work was supported by an MRC New Investigator Research Grant (grant number MR/R001898/1) awarded to FH and by a PhD studentship from the BBSRC Midlands Integrative Biosciences Training Partnership (MIBTP) awarded to NEH.

## Author Contributions

N.E.H. and F.H. designed research; N.E.H and J.L.L. performed research; N.E.H. analyzed data; N.E.H. wrote the paper; N.E.H, J.L.L and F.H. edited the paper.

