## Supplementary Information for "Transcriptome analysis of *Pseudomonas aeruginosa* biofilm infection in an *ex vivo* pig model of the cystic fibrosis lung"

*SUPPLEMENTARY FIGURES AND TABLES*

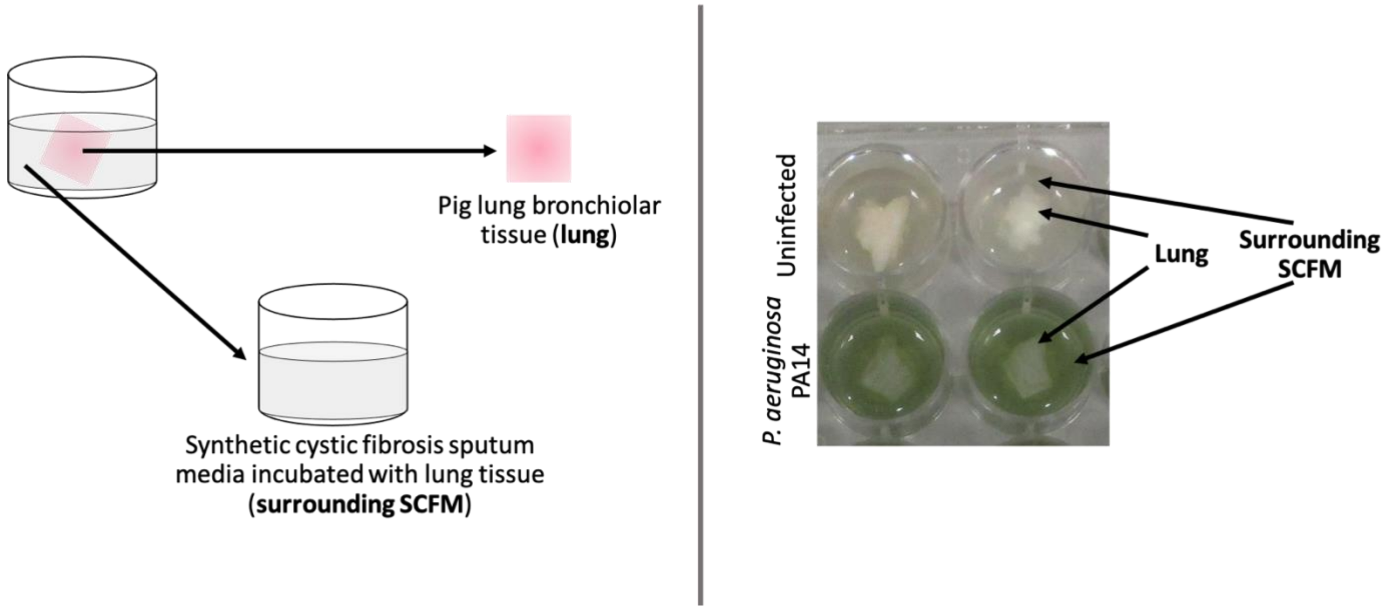


**Figure S1.** A diagram and photograph demonstrating the two environments of the *ex vivo* pig lung (EVPL) model from which *Pseudomonas aeruginosa* PA14 RNA was extracted and sequenced: the lung tissue and surrounding synthetic cystic fibrosis sputum media (SCFM). A 400 µl SCFM-agarose pad was made in each well of 24-well plates, and then the EVPL bronchiolar tissue was dissected into ~5 mm x 5 mm squares. The lung tissue pieces were placed on top of the SCFM-agarose pad and infected with PA14, with uninfected tissue used as a control. Each lung piece was then surrounded by 500 µl SCFM (surrounding SCFM) and the plate covered with a Breathe-Easier® membrane (Diversified Biotech), and incubated at 37 °C for the desired length of time. The surrounding SCFM was removed from the well for RNA extraction and the lung tissue piece bead beaten to retrieve the associated biofilm for RNA extraction.

**Table S1.** The concentration of RNA (ng µl$^{-1}$) extracted from 6 replicate *in vitro* synthetic cystic fibrosis sputum media (SCFM) *Pseudomonas aeruginosa* PA14 cultures following 7 d incubation at 37 °C. The concentration of RNA extracted from pig lung tissue-associated biofilm 7 d post infection (PI) with PA14 is also shown, from two independent lungs. Each row is an individual sample. Concentrations were measured using the Qubit™ RNA BR (broad range) Assay kit. The total RNA was determined from a final elution volume of 47 µl.

| **Environment** | **RNA concentration (ng µl$^{-1}$)** | **Total RNA (ng)** |
|---|---|---|
| SCFM | 0 | 0 |
| SCFM | 0 | 0 |
| SCFM | 36.4 | 1710.8 |
| SCFM | 24.4 | 1146.8 |
| SCFM | 0 | 0 |
| SCFM | 0 | 0 |
| Lung 1 | 27.4 | 1287.8 |
| Lung 1 | 33.8 | 1588.6 |
| Lung 1 | 25.8 | 1212.6 |
| Lung 2 | 51.6 | 2425.2 |
| Lung 2 | 14.2 | 667.4 |
| Lung 2 | 60.2 | 2829.4 |

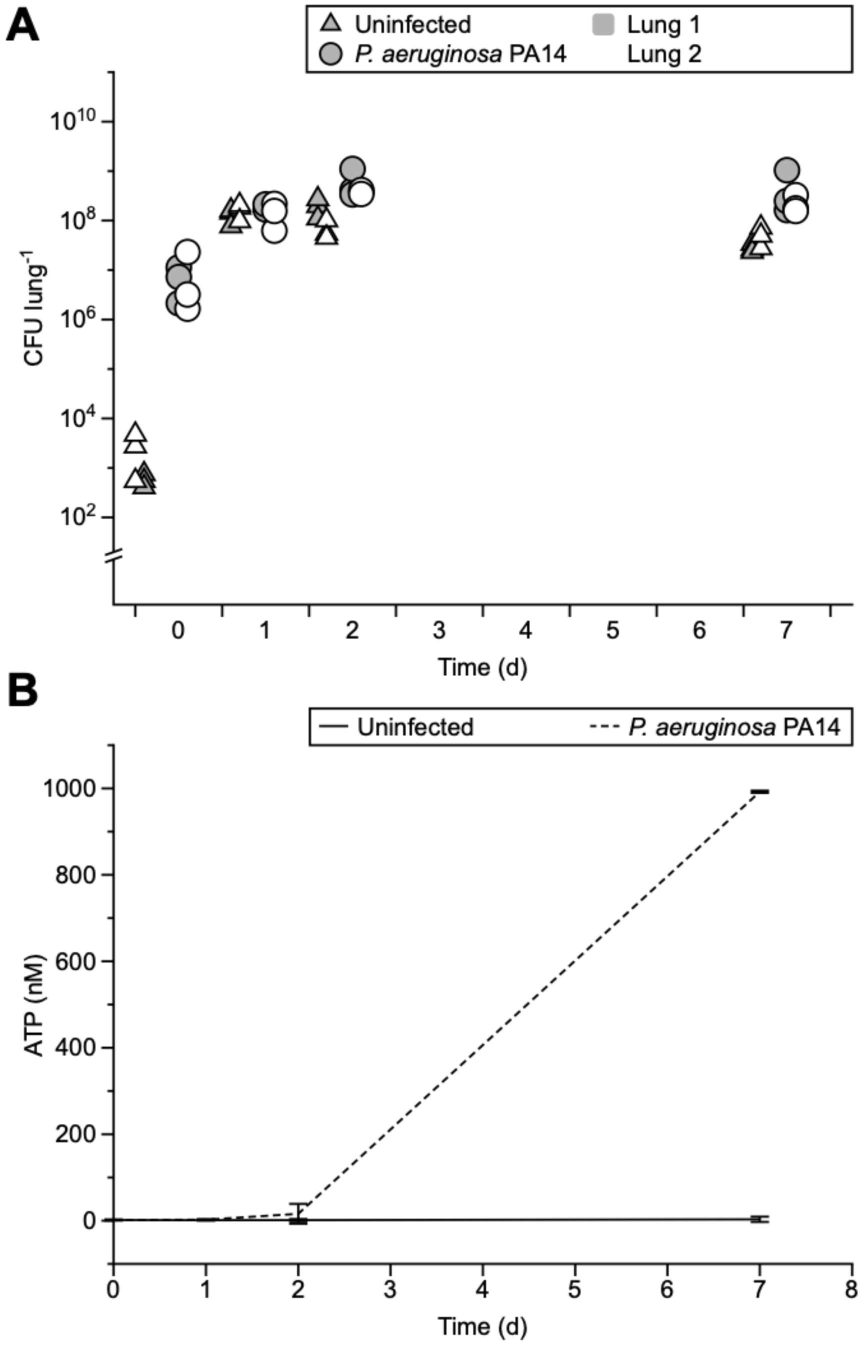


**Figure S2.** *Ex vivo* pig lung (EVPL) tissue from two independent lungs were infected with *Pseudomonas aeruginosa* PA14, with uninfected tissue as a control, for 7 d. The biofilm was recovered at 0 d, 1 d, 2 d and 7 d post infection (PI). **(A)** The colony forming units (CFU) per lung piece for three replicate lung pieces from each lung (lung 1 = filled, lung 2 = unfilled) per infection condition, determined at each time point. The uninfected (triangles) tissue CFU is of an undetermined endogenous species that was not detectable from *P. aeruginosa* PA14 infected tissue. **(B)** The ATP (nM) present in the recovered biofilms at each time point as a measure of microbial cell viability determined by a BacTiter-Glo™ microbial cell viability assay. The graph shows the average ATP determined from three replicate lung pieces from two independent lungs (6 repeats per time point and condition in total), error bars represent the standard deviation.

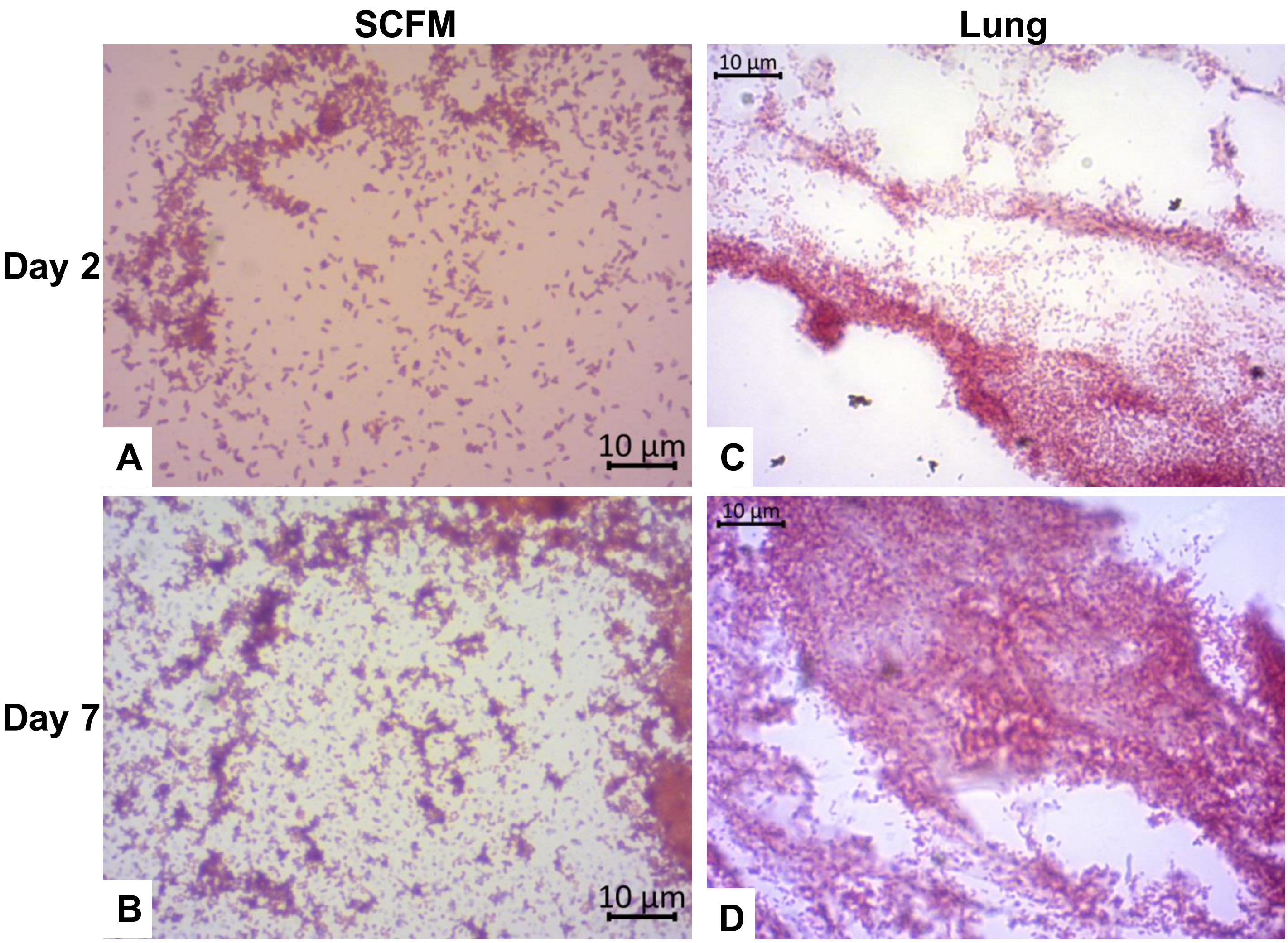


**Figure S3.** Micrographs of Gram-stained *Pseudomonas aeruginosa* PA14 grown in **(A,B)** synthetic cystic fibrosis sputum media (SCFM) and on **(C,D)** *ex vivo* pig lung (EVPL) tissue for 2 d and 7 d. EVPL images as used in [13]. All images are x100 magnification.

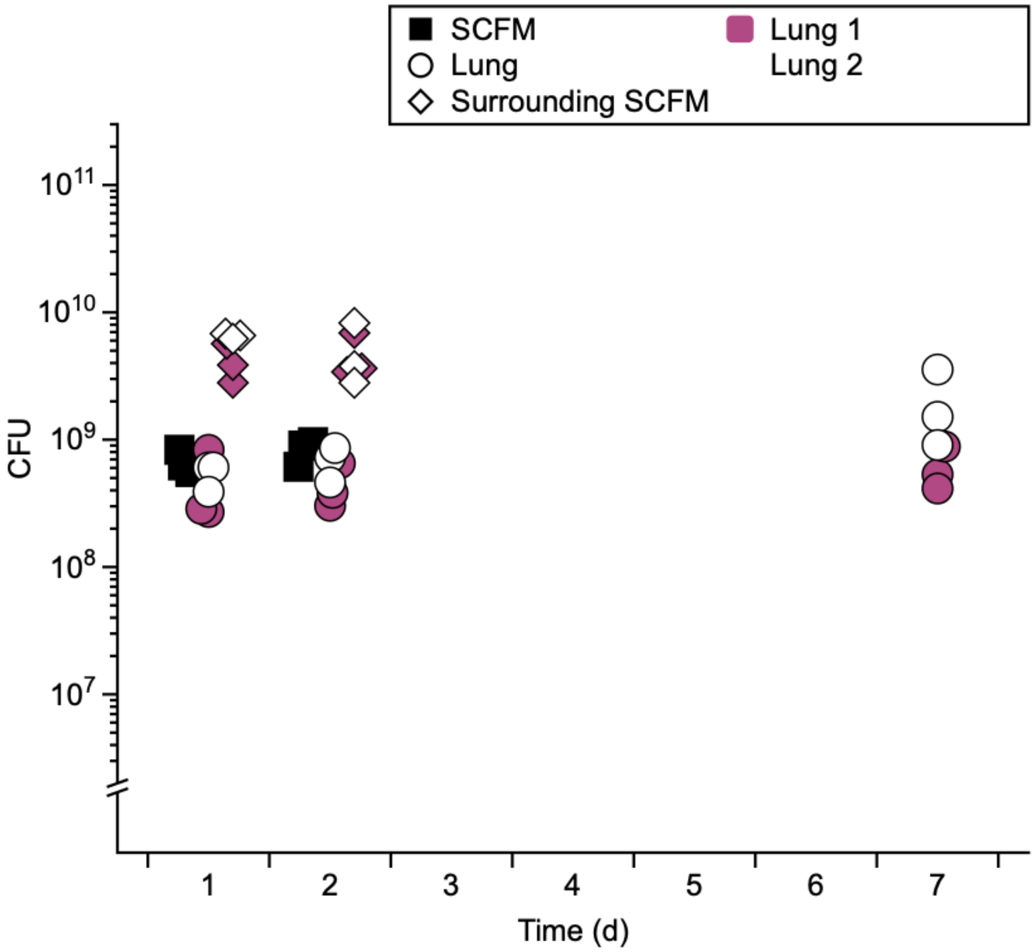


**Figure S4.** *Pseudomonas aeruginosa* PA14 colony forming units (CFU) of samples set up simultaneously to samples from which RNA was extracted. Three *ex vivo* pig lung (EVPL) tissue pieces from two independent lungs per time point were infected with PA14 (lung) and surrounded with synthetic cystic fibrosis sputum media (SCFM) (surrounding SCFM). Three replicate 1 ml *in vitro* SCFM PA14 cultures were also set up per time point (SCFM), and all samples were incubated at 37 °C. Destructive sampling was performed at 1 d, 2 d and 7 d post infection. The CFU per lung, and CFU $ml^{-1}$ for all SCFM samples, was determined. The y-axis is log scale. CFU in the surrounding SCFM at 1 d and 2 d was approximately one $log_{10}$ higher than CFU in *in vitro* SCFM and in lung tissue-associated biofilm.

**Table S2.** Percentage of reads aligned to the pig (*Sus scrofa*) genome and *Pseudomonas aeruginosa* PA14 genome from each sample. Each row shows an individual sample. The reads from the lung-associated biofilm and surrounding synthetic cystic fibrosis sputum media (SCFM) samples, from two independent lungs, were aligned to the *S. scrofa* genome and any reads that aligned were removed. The remaining reads and *in vitro* SCFM samples were then aligned to the PA14 genome.

| Environment | Time (d) | % Reads aligned to pig | % Reads aligned to PA14 |
|---|---|---|---|
| SCFM | 1 | | 95.30 |
| SCFM | 1 | | 98.39 |
| SCFM | 1 | | 99.30 |
| SCFM | 2 | | 99.58 |
| SCFM | 2 | | 98.88 |
| SCFM | 2 | | 99.36 |
| Lung 1 | 1 | 0.32 | 98.30 |
| Lung 1 | 1 | 0.23 | 99.76 |
| Lung 1 | 1 | 0.19 | 99.46 |
| Lung 2 | 1 | 0.37 | 96.17 |
| Lung 2 | 1 | 0.25 | 93.19 |
| Lung 2 | 1 | 0.35 | 95.34 |
| Lung 1 | 2 | 0.08 | 99.47 |
| Lung 1 | 2 | 0.08 | 99.46 |
| Lung 1 | 2 | 0.09 | 98.83 |
| Lung 2 | 2 | 0.07 | 98.17 |
| Lung 2 | 2 | 0.03 | 99.25 |
| Lung 2 | 2 | 0.19 | 96.86 |
| Lung 1 | 7 | 0.06 | 99.70 |
| Lung 1 | 7 | 0.08 | 99.13 |
| Lung 1 | 7 | 0.07 | 98.28 |
| Lung 2 | 7 | 0.06 | 98.65 |
| Lung 2 | 7 | 0.06 | 99.52 |
| Lung 2 | 7 | 0.02 | 99.10 |
| Surrounding SCFM (Lung 1) | 1 | 0.12 | 99.39 |
| Surrounding SCFM (Lung 1) | 1 | 0.87 | 97.56 |
| Surrounding SCFM (Lung 1) | 1 | 0.39 | 98.57 |
| Surrounding SCFM (Lung 2) | 1 | 0.27 | 97.57 |
| Surrounding SCFM (Lung 2) | 1 | 0.16 | 97.84 |
| Surrounding SCFM (Lung 2) | 1 | 1.04 | 94.61 |
| Surrounding SCFM (Lung 1) | 2 | 0.02 | 99.56 |
| Surrounding SCFM (Lung 1) | 2 | 0.08 | 99.27 |
| Surrounding SCFM (Lung 1) | 2 | 0.02 | 99.61 |
| Surrounding SCFM (Lung 2) | 2 | 0.08 | 98.71 |
| Surrounding SCFM (Lung 2) | 2 | 0.09 | 99.06 |

| | | | |
|---|---|---|---|
| Surrounding SCFM (Lung 2) | 2 | 0.10 | 98.92 |

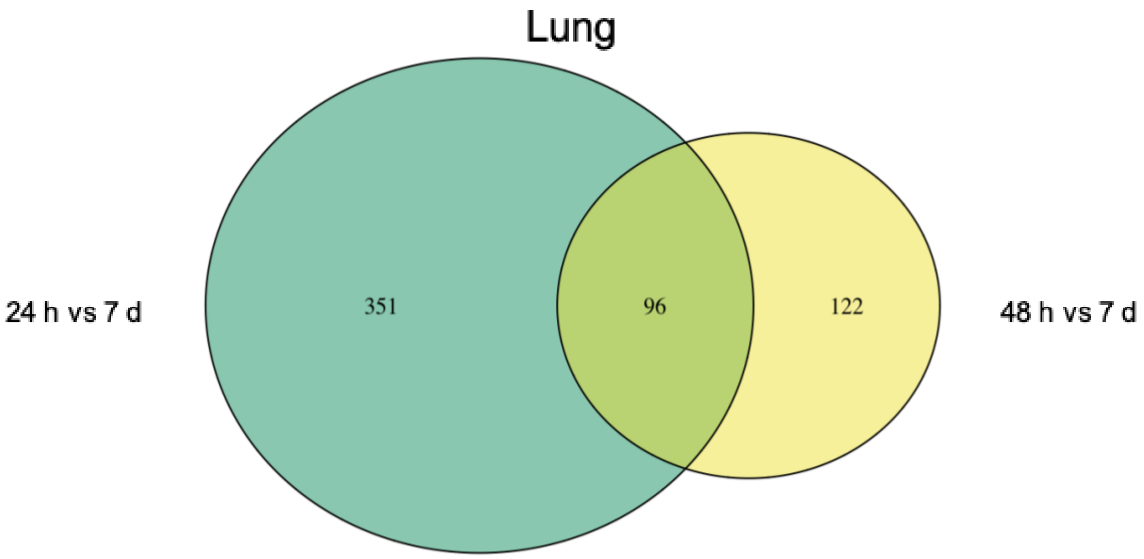


**Figure S5.** Venn diagram of the number of significant differentially expressed *Pseudomonas aeruginosa* PA14 genes (DEGs) from each contrast of interest, using threshold values of $P < 0.05$ and $\log_2$ fold change ≥ |1.5|. The contrasts shown are all of samples from the *ex vivo* pig lung tissue-associated *Pseudomonas aeruginosa* PA14 biofilm. Expression at 24 h post infection (PI) was compared with 7 d PI (green, left), and 48 h PI compared with 7 d PI (yellow, right). The shared DEGs between each contrast are genes that are either underexpressed or overexpressed in both comparisons. Genes that were significant DEGs in both contrasts at each time point, but in opposite directions, were not considered to be shared.

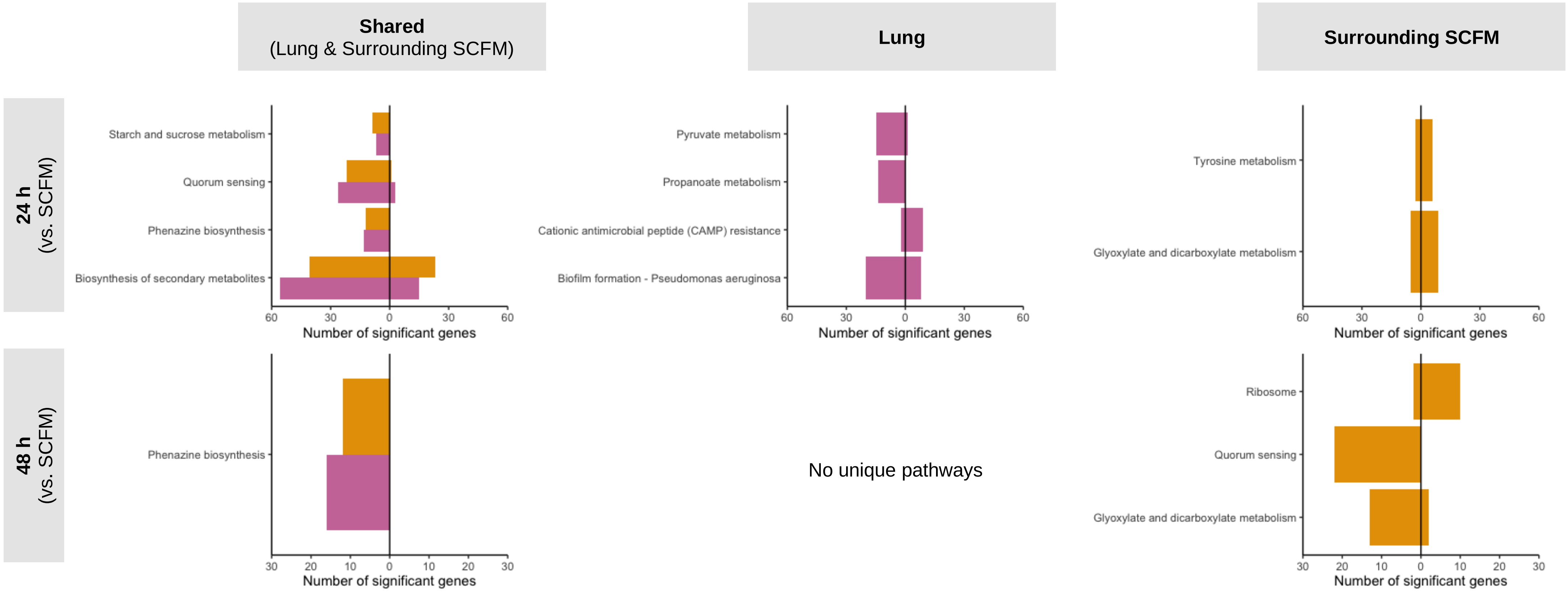


**Figure S6.** Significantly enriched ($P < 0.05$) *Pseudomonas aeruginosa* PA14 KEGG pathways in the *ex vivo* pig lung (EVPL) tissue-associated biofilm (lung) or surrounding synthetic cystic fibrosis sputum media (SCFM) (surrounding SCFM) compared with *in vitro* SCFM growth (SCFM), 24 h and 48 h post infection (PI). The bar charts show the number of significant differentially expressed genes ($P < 0.05$ and $\log_2$ fold change $\geq |1.5|$) from each pathway. Genes to the left of the vertical line (x = 0) are underexpressed in the lung and surrounding SCFM compared with *in vitro* SCFM, and genes to the right of the line are overexpressed. The shared graphs show pathways that are significantly enriched in both contrasts at that time point (lung-associated biofilm versus SCFM and surrounding SCFM versus SCFM) and the individual graphs show pathways only significant for that comparison. The lung-associated biofilm genes are shown by the purple bars (bottom bars in shared graphs) and the surrounding SCFM in the orange bars (top bars in shared graphs).

**Table S3.** All genes associated with the *Pseudomonas aeruginosa* PA14 KEGG pathway quorum sensing as listed on Pseudomonas.com [23]. Full table of results from which Table 1 was taken. PA14 expression in the two locations of the *ex vivo* pig lung model: the lung tissue-associated biofilm (lung) and the synthetic cystic fibrosis sputum media (SCFM) surrounding the lung tissue (surrounding SCFM), was compared with *in vitro* SCFM (SCFM) growth. Samples were compared at 24 h and 48 h post infection. The * denotes the only contrast where the quorum sensing KEGG pathway was not significantly enriched.

| **Locus tag** | **Gene name** | **Gene product** | **24 h: Lung vs SCFM** | **48 h: Lung vs SCFM*** | **24 h: Surrounding SCFM vs SCFM** | **48 h: Surrounding SCFM vs SCFM** |
|---|---|---|---|---|---|---|
| PA14_02720 | | Hypothetical protein | -1.16 | 0.87 | -1.51 | 0.12 |
| PA14_04900 | *ftsY* | Signal recognition particle receptor FtsY | 0.93 | 0.50 | 0.00 | 0.48 |
| PA14_05550 | *oprM* | Major intrinsic multiple antibiotic resistance efflux outer membrane protein OprM precursor | 1.00 | -0.33 | 0.87 | 0.57 |
| PA14_07850 | | ABC transporter substrate-binding protein | 0.71 | 0.91 | 0.43 | 0.14 |
| PA14_07860 | | ABC transporter ATP-binding protein | 1.09 | -0.11 | 1.40 | -0.51 |
| PA14_07870 | | ABC transporter substrate-binding protein | 0.38 | -1.27 | 1.41 | -0.96 |
| PA14_07890 | | ABC transporter permease | -0.33 | -0.80 | 0.73 | -0.97 |
| PA14_07900 | | ABC transporter permease | -0.29 | -0.64 | 0.55 | -0.86 |
| PA14_07940 | *trpE* | Anthranilate synthase component I | 0.62 | 0.47 | 0.22 | -0.09 |
| PA14_08340 | *trpG* | Anthranilate synthase component II | 0.64 | 0.39 | 0.26 | 0.08 |
| PA14_08370 | *vfr* | cAMP-regulatory protein | 1.25 | 0.70 | 0.79 | 0.62 |
| PA14_08695 | *secE* | Preprotein translocase subunit SecE | -0.83 | 0.55 | -1.31 | -0.14 |
| PA14_09050 | *secY* | Preprotein translocase subunit SecY | 0.75 | 1.94 | -0.32 | 1.07 |
| PA14_09410 | *phzG1* | Pyrodoxamine 5'-phosphate oxidase | -3.90 | -4.38 | -1.97 | -2.55 |
| PA14_09420 | *phzF1* | Phenazine biosynthesis protein | -5.48 | -4.90 | -2.98 | -3.07 |
| PA14_09440 | *phzE1* | Phenazine biosynthesis protein PhzE | -5.82 | -5.04 | -3.26 | -3.33 |

| | | | | | | |
|---|---|---|---|---|---|---|
| PA14_09450 | *phzD1* | Phenazine biosynthesis protein PhzD | -6.09 | -5.51 | -3.17 | -3.09 |
| PA14_09460 | *phzC1* | Phenazine biosynthesis protein PhzC | -4.82 | -5.15 | -1.79 | -2.92 |
| PA14_09470 | *phzB1* | Phenazine biosynthesis protein | -4.50 | -4.19 | -1.37 | -2.93 |
| PA14_09520 | *mexI* | RND efflux transporter | -1.97 | -2.83 | -1.73 | -1.53 |
| PA14_09530 | *mexH* | RND efflux membrane fusion protein | -0.85 | -2.74 | -0.37 | -0.93 |
| PA14_11400 | *ribD* | Riboflavin-specific deaminase/reductase | 1.01 | -0.72 | 0.58 | 0.09 |
| PA14_11510 | *ribA* | GTP cyclohydrolase II | 0.54 | 0.05 | 0.08 | 0.27 |
| PA14_14610 | *yajC* | Preprotein translocase subunit YajC | 0.95 | 0.45 | 0.59 | 1.05 |
| PA14_15520 | *trbJ* | Conjugal transfer protein TrbJ | 0.64 | 0.76 | 0.28 | -0.11 |
| PA14_15540 | | Mating pair formation protein TrbL | 0.30 | 0.15 | 0.10 | -0.33 |
| PA14_15960 | *ffh* | Signal recognition particle protein Ffh | 0.01 | 0.49 | -0.79 | 0.16 |
| PA14_16250 | *lasB* | Elastase LasB | -3.35 | -3.01 | -2.28 | -1.62 |
| PA14_17140 | | Membrane-associated zinc metalloprotease | 1.08 | 1.94 | -0.44 | 0.69 |
| PA14_19100 | *rhlA* | Rhamnosyltransferase chain A | -1.21 | -0.35 | -1.17 | -0.49 |
| PA14_19110 | *rhlB* | Rhamnosyltransferase chain B | -1.79 | -1.33 | -1.08 | -0.90 |
| PA14_19120 | *rhlR* | Transcriptional regulator RhlR | -0.77 | -1.23 | -0.06 | -0.70 |
| PA14_19130 | *rhlI* | Autoinducer synthesis protein RhlI | -0.27 | 0.17 | -0.90 | -0.55 |
| PA14_20580 | *amiC* | Aliphatic amidase expression-regulating protein | -0.47 | 0.90 | -0.63 | -0.02 |
| PA14_20610 | *lecB* | Fucose-binding lectin PA-IIL | -3.43 | -2.90 | -2.52 | -1.95 |
| PA14_21090 | | Hypothetical protein | -2.20 | 0.16 | -1.07 | 0.48 |
| PA14_21110 | *plcN* | Non-hemolytic phospholipase C | -0.86 | 1.16 | -1.07 | 0.05 |
| PA14_21340 | *fadD2* | Long-chain-fatty-acid--CoA ligase | -0.14 | -0.19 | 0.16 | -0.25 |

| | | | | | | |
|---|---|---|---|---|---|---|
| PA14_21370 | *fadD1* | Long-chain-fatty-acid--CoA ligase | 0.30 | -0.20 | -0.97 | -0.47 |
| PA14_21910 | | ABC transporter ATP-binding protein | -0.06 | -0.74 | 0.31 | -0.59 |
| PA14_21920 | | ABC transporter permease | 0.23 | 0.07 | 0.28 | -0.62 |
| PA14_21930 | | ABC transporter permease | -0.07 | 0.80 | -0.15 | -0.47 |
| PA14_21960 | | Hypothetical protein | 0.25 | -0.74 | 0.41 | -0.35 |
| PA14_25980 | *aroF* | Phospho-2-dehydro-3-deoxyheptonate aldolase | 0.61 | 0.14 | 0.03 | 0.57 |
| PA14_27330 | | Phospho-2-dehydro-3-deoxyheptonate aldolase | 0.13 | -3.18 | 0.52 | -2.06 |
| PA14_30630 | *pqsH* | FAD-dependent monooxygenase | -0.58 | -1.31 | 0.38 | -0.43 |
| PA14_30850 | | TrbI-like protein | 0.50 | 1.81 | -1.77 | 0.16 |
| PA14_30860 | | TrbG-like protein | -0.04 | 1.72 | -0.40 | -0.66 |
| PA14_30870 | | Conjugal transfer protein TrbF | -0.52 | -0.31 | 0.04 | -0.80 |
| PA14_30880 | | Conjugal transfer protein TrbL | -0.90 | 1.31 | -1.15 | -0.31 |
| PA14_30900 | | Conjugal transfer protein TrbJ | -1.14 | -0.18 | -0.72 | -1.49 |
| PA14_30910 | | Conjugal transfer ATPase TrbE | -1.02 | 0.36 | -0.97 | -0.52 |
| PA14_30930 | | TrbC-like protein | -0.80 | 1.38 | -1.39 | -0.97 |
| PA14_30940 | | Conjugal transfer protein | -1.04 | 0.66 | -1.35 | -0.53 |
| PA14_31290 | *pa1L* | PA-I galactophilic lectin | -3.94 | 1.52 | -3.73 | 0.04 |
| PA14_31300 | | Hypothetical protein | -1.34 | 0.03 | -0.11 | -0.04 |
| PA14_39880 | *phzG2* | Pyridoxamine 5'-phosphate oxidase | -5.51 | -5.15 | -2.91 | -3.14 |
| PA14_39890 | *phzF2* | Phenazine biosynthesis protein | -5.48 | -4.86 | -2.96 | -3.09 |
| PA14_39945 | *phzC2* | Phenazine biosynthesis protein PhzC | -5.89 | -5.68 | -2.95 | -3.75 |
| PA14_39960 | *phzB2* | Phenazine biosynthesis protein | -5.24 | -5.09 | -2.68 | -3.97 |
| PA14_39970 | *phzA2* | Phenazine biosynthesis protein | -4.93 | -4.40 | -3.92 | -4.52 |
| PA14_40260 | | Hypothetical protein | -2.17 | -3.26 | -0.17 | -0.88 |
| PA14_40290 | *lasA* | LasA protease | -2.09 | -2.22 | -1.36 | -1.53 |
| PA14_41920 | *aroF-1* | Phospho-2-dehydro-3-deoxyheptonate aldolase | -0.26 | -0.15 | -0.48 | -0.43 |

| | | | | | | |
|---|---|---|---|---|---|---|
| PA14_43340 | *kdpE* | Two-component response regulator KdpE | -0.34 | -1.26 | -0.12 | -0.56 |
| PA14_45940 | *lasI* | Autoinducer synthesis protein LasI | -0.10 | -0.57 | 0.94 | -1.10 |
| PA14_45960 | *lasR* | Transcriptional regulator LasR | 0.03 | -0.53 | 0.82 | 0.03 |
| PA14_47370 | | Signal peptidase | 0.62 | 0.73 | 0.34 | 0.60 |
| PA14_49760 | *rhlC* | Rhamnosyltransferase 2 | -1.71 | -1.61 | -0.96 | -1.01 |
| PA14_50520 | *braC* | Branched-chain amino acid transport protein BraC | 0.70 | -0.68 | 1.01 | -0.69 |
| PA14_50530 | *braD* | Branched-chain amino acid transport protein BraD | 0.19 | -0.56 | -0.04 | -1.16 |
| PA14_50540 | *livM* | Leucine/isoleucine/valine transporter permease subunit | 0.07 | -1.26 | -0.19 | -1.65 |
| PA14_50550 | *livG* | Leucine/isoleucine/valine transporter ATP-binding subunit | -0.37 | -1.37 | -0.23 | -1.65 |
| PA14_50560 | *braG* | Branched-chain amino acid transport protein BraG | -0.39 | -1.24 | -0.45 | -1.90 |
| PA14_51340 | *mvfR* | Transcriptional regulator MvfR | -0.44 | -0.09 | -0.83 | -0.22 |
| PA14_51350 | *phnB* | Anthranilate synthase component II | -0.56 | -1.36 | -0.75 | -0.30 |
| PA14_51360 | *phnA* | Anthranilate synthase component I | -0.78 | -1.20 | -1.07 | -0.31 |
| PA14_51380 | *pqsE* | Quinolone signal response protein | -0.82 | -0.57 | -1.29 | 0.12 |
| PA14_51390 | *pqsD* | 3-oxoacyl-ACP synthase | -0.99 | -1.83 | -0.97 | -0.47 |
| PA14_51410 | *pqsC* | PqsC | -1.32 | -1.87 | -0.89 | -0.55 |
| PA14_51420 | *pqsB* | PqsB | -1.02 | -0.86 | -0.96 | 0.18 |
| PA14_51430 | *pqsA* | PqsA | -1.06 | -0.97 | -1.34 | -0.20 |
| PA14_53360 | *plcH* | Hemolytic phospholipase C | -0.16 | 1.50 | -0.53 | 0.54 |
| PA14_54350 | *lepB* | Signal peptidase I | 0.20 | 0.07 | -0.43 | 0.17 |
| PA14_57220 | *secA* | Preprotein translocase subunit SecA | 0.38 | 0.83 | 0.01 | 0.52 |
| PA14_58360 | *dppA2* | Dipeptide ABC transporter substrate-binding protein DppA2 | 2.25 | 2.33 | 0.80 | 0.44 |
| PA14_62810 | *secG* | Preprotein translocase subunit SecG | 0.24 | 2.50 | -0.88 | 0.76 |

| | | | | | | |
|---|---|---|---|---|---|---|
| PA14_63150 | *pmrA* | Two-component response regulator | 1.51 | 0.37 | 0.14 | -0.28 |
| PA14_63160 | *pmrB* | PmrB: two-component regulator system signal sensor kinase PmrB | 1.40 | 0.10 | 0.27 | -0.35 |
| PA14_63480 | | Amino acid permease | 0.06 | 1.18 | 0.06 | 0.08 |
| PA14_63650 | | Hypothetical protein | 0.63 | -0.09 | -0.33 | 0.79 |
| PA14_64860 | | ABC transporter ATP-binding protein | -2.97 | -1.62 | -2.12 | -0.91 |
| PA14_64870 | | ABC transporter ATP-binding protein | -2.29 | -1.11 | -1.71 | -0.89 |
| PA14_64880 | | Branched chain amino acid ABC transporter permease | -2.04 | -0.39 | -1.82 | -0.60 |
| PA14_64890 | | Branched chain amino acid ABC transporter permease | -0.80 | -0.09 | -0.28 | -0.03 |
| PA14_64900 | | ABC transporter substrate-binding protein | -0.98 | -0.82 | 0.01 | -0.32 |
| PA14_65310 | *hfq* | RNA-binding protein Hfq | 1.59 | 0.52 | 1.87 | 0.45 |
| PA14_67720 | *secB* | Preprotein translocase subunit SecB | -0.10 | -1.00 | 0.40 | -0.19 |
| PA14_70200 | *dppA5* | Dipeptide ABC transporter substrate-binding protein DppA5 | 0.38 | 0.28 | 0.37 | -0.24 |
| PA14_72560 | *np20* | Transcriptional regulator np20 | -1.32 | -2.17 | 0.14 | -0.31 |
| PA14_73410 | | Inner membrane protein translocase component YidC | 0.03 | 0.75 | -1.23 | 0.40 |

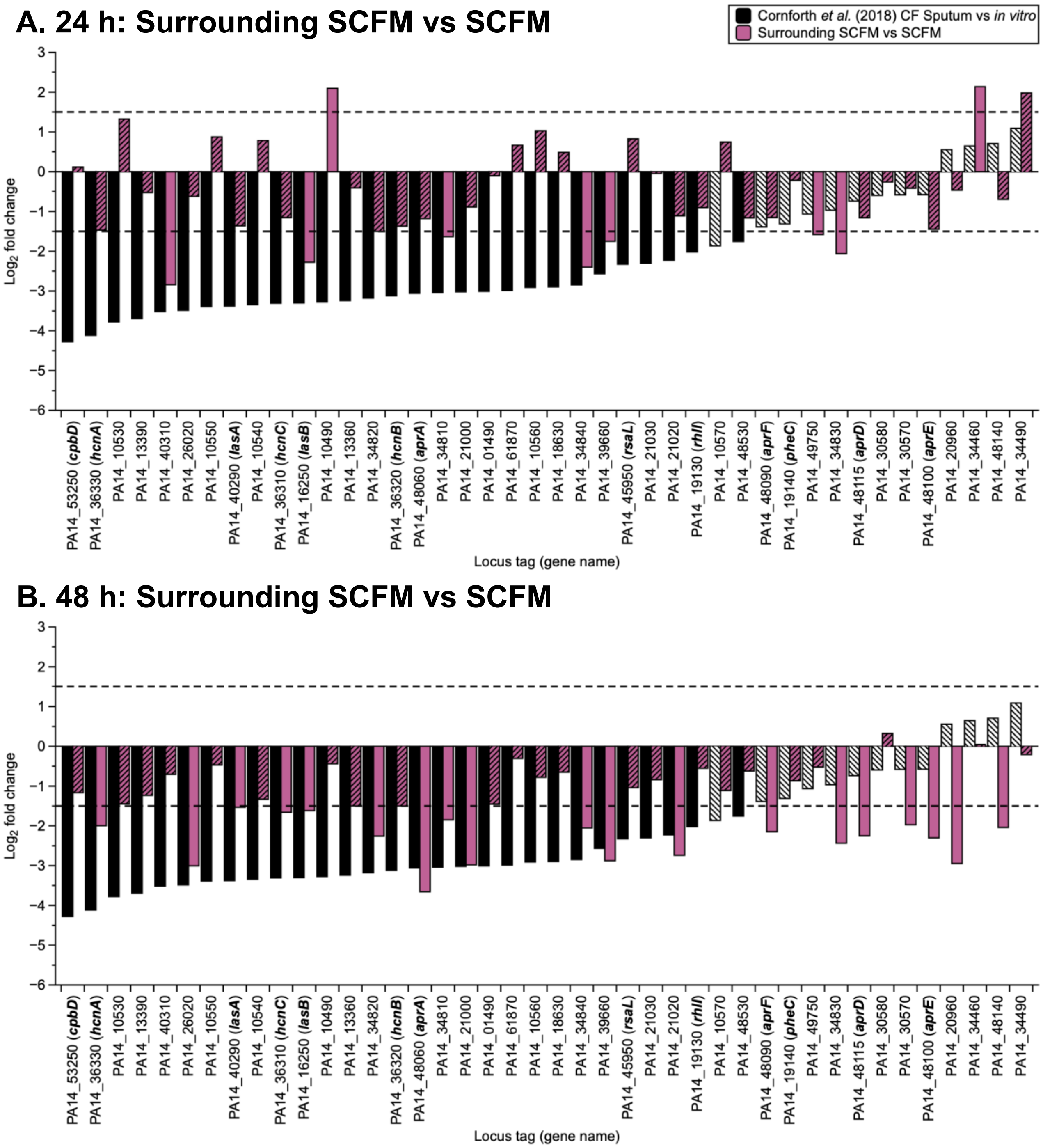


**Figure S7.** The $\log_2$ fold change (lfc) in expression of 42 *Pseudomonas aeruginosa* quorum sensing genes, controlled by the las regulon, conserved in human infection. The expression of *P. aeruginosa* PA14 grown in the synthetic cystic fibrosis sputum media (SCFM) surrounding *ex vivo* pig lung tissue pieces (surrounding SCFM) versus *in vitro* SCFM at 24 h (**A**) and 48 h (**B**) post infection are shown by the purple bars. Each graph also includes expression of the gene set by *P. aeruginosa* from human cystic fibrosis sputum versus *in vitro* conditions taken from Cornforth *et al.* [7], shown by the black bars. The locus tags shown are for *P. aeruginosa* PA14 with gene names in bold where appropriate. Bars with the striped fill are not significantly differentially expressed for that contrast ($P < 0.05$, lfc $\geq |1.5|$). The dashed lines represent the threshold lfc value for differential expression used by both studies.

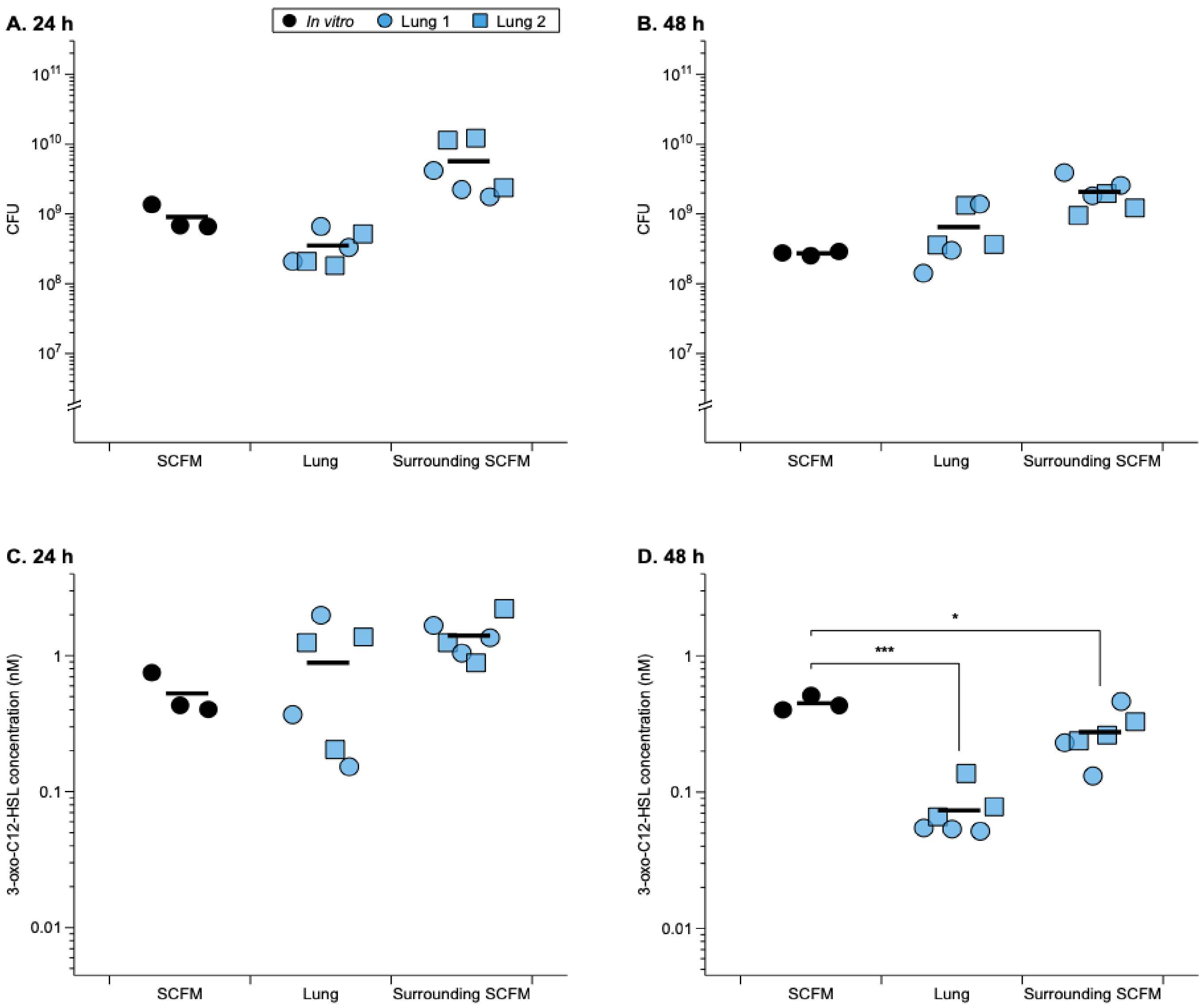


**Figure S8.** *Pseudomonas aeruginosa* PA14 was grown in synthetic cystic fibrosis sputum media (SCFM) *in vitro*, and in the *ex vivo* pig lung model (EVPL) environments: lung-associated biofilm (lung) and the surrounding SCFM, for 24 h and 48 h at 37 °C. Bars denote the mean for each environment on each graph. **(A,B)** Aliquots of the SCFM cultures and lung homogenate were taken and the CFU $ml^{-1}$ and CFU per lung respectively determined. **(C,D)** The remaining SCFM cultures (*in vitro* SCFM and surrounding SCFM) and lung homogenate were filter sterilized. The amount of 3-oxo-C12-HSL present in each filtered supernatant was measured using the bioreporter *Escherichia coli* JM107 with a pSB1142 plasmid, and the concentration determined using a standard curve. Asterisks denote a significant difference in the amount of 3-oxo-C12-HSL between growth environments. ANOVA found no significant difference at 24 h ($F_{2,12} = 2.53$, $P$ = 0.12), but there was a significant difference at 48 h (ANOVA: F2,12 = 32.83, $P < 0.01$). Post-hoc analysis showed there was significantly more 3-oxo-C12-HSL produced in SCFM than the lung ($P$ 0.01) and surrounding SCFM ($P = 0.02$) at 48 h post infection.

**Table S4.** *Pseudomonas aeruginosa* PA14 expression of three genes associated with quorum quenching [36] in the two locations of the *ex vivo* pig lung model: the lung tissue-associated biofilm (lung) and the synthetic cystic fibrosis sputum media (SCFM) surrounding the lung tissue (surrounding SCFM), was compared with *in vitro* SCFM (SCFM) growth at 24 h and 48 h post infection. The $\log_2$ fold change (lfc) is shown, and those in bold and underlined were found to be significantly differentially expressed ($P < 0.05$).

| **Locus tag** | **Gene name** | **Gene product** | **24 h: Lung vs SCFM** | **48 h: Lung vs SCFM** | **24 h: Surrounding SCFM vs SCFM** | **48 h: Surrounding SCFM vs SCFM** |
|---|---|---|---|---|---|---|
| PA14_03980 | | Hypothetical protein | 0.41 | -0.48 | 0.52 | -0.35 |
| PA14_33820 | *pvdQ* | Penicillin acylase-related protein | -1.15 | <u>**-1.88**</u> | 0.11 | -0.39 |
| PA14_50980 | *pac* | Penicillin amidase | -0.09 | 0.04 | -0.15 | -0.29 |

**Table S5.** All 52 genes predicted to be involved in *Pseudomonas aeruginosa* PA14 antimicrobial resistance by the Comprehensive Antibiotic Resistance Database (CARD) [23]. PA14 expression in the two locations of the *ex vivo* pig lung model: the lung tissue-associated biofilm (lung) and the synthetic cystic fibrosis sputum media (SCFM) surrounding the lung tissue (surrounding SCFM), was compared with *in vitro* SCFM (SCFM) growth. Samples were compared at 24 h and 48 h post infection. The $\log_2$ fold change (lfc) is shown, and those in bold and underlined were found to be significantly differentially expressed ($P < 0.05$). The model name detailed by the CARD is also shown alongside the gene locus tag and gene product.

| **Locus tag** | **Model name** | **Gene product** | **24 h: Lung vs SCFM** | **48 h: Lung vs SCFM** | **24 h: Surrounding SCFM vs SCFM** | **48 h: Surrounding SCFM vs SCFM** |
|---|---|---|---|---|---|---|
| PA14_01940 | TriA | RND efflux membrane fusion protein | -0.04 | -0.90 | 0.08 | -0.88 |
| PA14_01960 | TriB | RND efflux membrane fusion protein | -0.55 | <u>**-1.72**</u> | -0.039 | -0.97 |
| PA14_01970 | TriC | RND efflux transporter | -0.56 | -1.10 | -0.24 | -0.86 |
| PA14_05520 | MexR | Multidrug resistance operon repressor MexR | 0.25 | 0.69 | -0.15 | 0.40 |
| PA14_05530 | MexA | RND multidrug efflux membrane fusion protein MexA | 0.48 | 0.66 | 0.24 | 0.69 |
| PA14_05540 | MexB | RND multidrug efflux transporter MexB | 0.95 | 0.94 | 0.28 | 0.99 |
| PA14_05550 | OprM | Major intrinsic multiple antibiotic resistance efflux outer membrane protein OprM precursor | 1.00 | -0.33 | 0.87 | 0.57 |
| PA14_09500 | OpmD | Outer membrane protein | <u>**-2.77**</u> | <u>**-3.41**</u> | <u>**-1.87**</u> | <u>**-1.79**</u> |
| PA14_09520 | MexI | RND efflux transporter | <u>**-1.97**</u> | <u>**-2.83**</u> | <u>**-1.73**</u> | <u>**-1.53**</u> |
| PA14_09530 | MexH | RND efflux membrane fusion protein | -0.85 | <u>**-2.74**</u> | -0.37 | -0.93 |
| PA14_09540 | MexG | Hypothetical protein | -0.60 | <u>**-2.32**</u> | -0.02 | -0.07 |
| PA14_10470 | bcr-1 | MFS transporter | <u>**2.06**</u> | -0.03 | 1.25 | -0.12 |
| PA14_10670 | APH(3')-IIb | Aminoglycoside 3'-phosphotransferase type IIB | -0.15 | 0.41 | -1.09 | 0.37 |
| PA14_10790 | PDC-9 | Beta-lactamase | 0.37 | 0.72 | 0.53 | 0.73 |
| PA14_16280 | nalC | Transcriptional regulator | -0.86 | -0.57 | 0.88 | 0.73 |
| PA14_16300 | ArmR | Hypothetical protein | <u>**-1.52**</u> | -0.80 | 1.00 | 1.16 |
| PA14_16790 | MexL | TetR family transcriptional regulator | -0.67 | <u>**-1.62**</u> | 0.25 | -0.14 |
| PA14_16800 | MexJ | Efflux transmembrane protein | 0.85 | 1.31 | 0.01 | 0.41 |

| | | | | | | |
|---|---|---|---|---|---|---|
| PA14_16820 | MexK | Efflux transmembrane protein | 1.13 | 1.29 | -0.36 | -0.01 |
| PA14_18080 | nalD | TetR family transcriptional regulator | 1.17 | 1.45 | **2.22** | **2.04** |
| PA14_18350 | arnA | Bifunctional UDP-glucuronic acid decarboxylase/UDP-4-amino-4-deoxy-L-arabinose formyltransferase | **2.34** | 1.42 | -0.39 | 0.25 |
| PA14_18760 | mexP | RND efflux membrane fusion protein | 0.91 | **1.84** | 1.14 | **1.76** |
| PA14_18780 | mexQ | RND efflux transporter | 0.84 | 1.44 | 0.70 | 1.19 |
| PA14_18790 | opmE | Outer membrane efflux protein | 0.88 | 1.27 | 0.72 | **1.61** |
| PA14_22760 | *Pseudomonas aeruginosa* CpxR | Two-component response regulator | 0.68 | -0.19 | 0.29 | -0.26 |
| PA14_31870 | MuxA | RND efflux membrane fusion protein | 1.17 | **1.67** | -0.01 | 0.43 |
| PA14_31890 | MuxB | RND efflux transporter | 0.86 | 0.37 | 0.02 | -0.01 |
| PA14_31900 | MuxC | Efflux transporter | **1.55** | 0.84 | 0.19 | 0.19 |
| PA14_31920 | OpmB | Outer membrane protein | **1.68** | 0.25 | 0.87 | 0.29 |
| PA14_32380 | OprN | Multidrug efflux outer membrane protein OprN precursor | -0.28 | -0.56 | -0.53 | -0.73 |
| PA14_32390 | MexF | RND multidrug efflux transporter MexF | -0.30 | 0.69 | -1.12 | -0.23 |
| PA14_32400 | MexE | RND multidrug efflux membrane fusion protein MexE | -0.42 | 0.82 | -1.28 | 0.49 |
| PA14_32410 | MexT | Transcriptional regulator MexT | -0.05 | -0.46 | -0.04 | 0.05 |
| PA14_32420 | MexS | Oxidoreductase | -0.40 | -0.63 | -0.30 | -0.03 |
| PA14_35170 | *Pseudomonas aeruginosa* soxR | Redox-sensing activator of soxS | -1.17 | -0.91 | -0.31 | -0.05 |
| PA14_38380 | MexZ | Transcriptional regulator | -1.16 | 0.10 | -0.74 | 0.48 |
| PA14_45890 | mexN | RND efflux transporter | -0.18 | 0.72 | -0.25 | 0.81 |
| PA14_45910 | mexM | RND efflux membrane fusion protein | 0.42 | **1.72** | -0.93 | 0.12 |
| PA14_46680 | PmpM | Transporter | 0.64 | 0.87 | -0.40 | 0.29 |
| PA14_49780 | FosA | Fosfomycin resistance protein | **-1.61** | -0.18 | -0.88 | -0.14 |
| PA14_55170 | *Pseudomonas aeruginosa* catB7 | Chloramphenicol acetyltransferase | -0.65 | -0.09 | -1.09 | -0.27 |

| | | | | | | |
|---|---|---|---|---|---|---|
| PA14_56880 | MexV | Membrane fusion protein | 0.70 | -0.21 | 0.23 | -0.65 |
| PA14_56890 | MexW | Multidrug efflux protein | -0.24 | -0.29 | -0.17 | -1.17 |
| PA14_59160 | CrpP | CrpP | -0.23 | **1.68** | -0.31 | 1.18 |
| PA14_60820 | OprJ | Outer membrane protein OprJ | 0.80 | 0.31 | **1.55** | 1.08 |
| PA14_60830 | MexD | Multidrug efflux RND transporter MexD | 0.86 | **1.61** | 0.25 | **1.58** |
| PA14_60850 | MexC | Multidrug efflux RND membrane fusion protein | 0.54 | **2.09** | -0.16 | **2.25** |
| PA14_60860 | Type B NfxB | Transcriptional regulator NfxB | -0.66 | -0.33 | -0.31 | 0.44 |
| PA14_63160 | basS | PmrB: two-component regulator system signal sensor kinase PmrB | 1.40 | 0.10 | 0.27 | -0.35 |
| PA14_65750 | OpmH | Outer membrane efflux protein | 0.06 | **-1.57** | 0.31 | -0.68 |
| PA14_65990 | *Pseudomonas aeruginosa* emrE | SMR multidrug efflux transporter | -0.45 | 1.11 | -0.39 | 0.16 |
| PA14_72760 | OXA-50 | Beta-lactamase | -0.30 | -0.30 | 0.15 | -0.16 |

**Table S6.** Minimum inhibitory concentration (MIC) of colistin and polymyxin B (μg $ml^{-1}$) for *Pseudomonas aeruginosa* PA14. The MIC value in cation-adjusted Mueller Hinton broth (CAMHB), the clinical standard, and synthetic cystic fibrosis sputum media (SCFM) was determined based on three replicates. The MIC values for the lung are from MIC tests performed using *P. aeruginosa* PA14 cells retrieved from the *ex vivo* pig lung tissue-associated biofilm. Infected lung tissue was bead beaten at 24 h and 48 h post infection. The MIC was determined for three replicate lung pieces from each of two independent lungs.

| **Antibiotic** | **CAMHB** | **SCFM** | **Lung (24 h)** | **Lung (48 h)** |
|---|---|---|---|---|
| Colistin | 2 | 2 | 8 | 16 |
| Polymyxin B | 1 | 2 | 8 | 4 |
